# Tumour mutations in long noncoding RNAs enhance cell fitness

**DOI:** 10.1101/2021.11.06.467555

**Authors:** Roberta Esposito, Andrés Lanzós, Taisia Polidori, Hugo Guillen-Ramirez, Bernard Mefi Merlin, Lia Mela, Eugenio Zoni, Isabel Büchi, Lusine Hovhannisyan, Finn McCluggage, Matúš Medo, Giulia Basile, Dominik F. Meise, Sunandini Ramnarayanan, Sandra Zwyssig, Corina Wenger, Kyriakos Schwarz, Adrienne Vancura, Núria Bosch-Guiteras, Marianna Kruithof-de Julio, Yitzhak Zimmer, Michaela Medová, Deborah Stroka, Archa Fox, Rory Johnson

## Abstract

Long noncoding RNAs (lncRNAs) can act as tumour suppressors or oncogenes to repress/promote tumour cell proliferation via RNA-dependent mechanisms. Recently, genome sequencing has identified elevated densities of tumour somatic single nucleotide variants (SNVs) in lncRNA genes. However, this has been attributed to phenotypically-neutral “passenger” processes, and the existence of positively-selected fitness-altering “driver” SNVs acting via lncRNAs has not been addressed. We developed and used *ExInAtor2*, an improved driver-discovery pipeline, to map pancancer and cancer-specific mutated lncRNAs across an extensive cohort of 2583 primary and 3527 metastatic tumours. The 54 resulting lncRNAs are mostly linked to cancer for the first time. Their significance is supported by a range of clinical and genomic evidence, and display oncogenic potential when experimentally expressed in matched tumour models. Our results revealed a striking SNV hotspot in the iconic *NEAT1* oncogene, which was ascribed by previous studies to passenger processes. To directly evaluate the functional significance of *NEAT1* SNVs, we used *in cellulo* mutagenesis to introduce tumour-like mutations in the gene and observed a consequent increase in cell proliferation in both transformed and normal backgrounds. Mechanistic analyses revealed that SNVs alter *NEAT1* ribonucleoprotein assembly and boost subnuclear paraspeckles. This is the first experimental evidence that mutated lncRNAs can contribute to the pathological fitness of tumour cells.

## Introduction

Tumours arise and develop via somatic mutations that confer a fitness advantage on cells ^1^. Such “driver” mutations exert their phenotypic effect by altering the function of genes or genomic elements, and are characterised by signatures of positive evolutionary selection ^2^. This is complicated by numerous “passenger” mutations, which do not impact cell phenotype and are evolutionarily neutral ^3^. Identification of driver mutations, and the “driver genes” through which they act, is a critical step towards understanding and treating cancer ^1,4^.

Most tumours are characterised by a limited and recurrent sequence of driver mutations, which promote disease hallmarks via functional changes to encoded oncogene or tumour suppressor proteins. However, the vast majority of somatic single nucleotide variants (SNVs) fall outside protein-coding genes ^5^. Combined with increasing awareness of the disease roles of noncoding genomic elements ^6^, this naturally raises the question of whether non-protein coding mutations can also shape cancer cell fitness ^7^. Growing numbers of both theoretical ^8–13^ and experimental studies ^2,14–17^ implicate noncoding SNVs in cell fitness by altering the function of elements such as enhancers, promoters, insulator elements and small RNAs ^18^.

Surprisingly, one important class of cancer-promoting noncoding genes has been largely overlooked: long noncoding RNAs (lncRNAs) ^19^. LncRNA transcripts are modular assemblages of functional elements that can interact with other nucleic acids and proteins via defined sequence or structural elements^20,21^. Of the >50,000 loci mapped in the human genome ^22^, hundreds of “cancer-lncRNAs” have been demonstrated to act as oncogenes or tumour suppressors ^23^. Their clinical importance is further supported by copy number variants (CNVs) ^24–26^, tumour-initiating transposon screens in mouse ^27^ and function-altering germline cancer variants ^28^.

We and others have previously reported statistical evidence for positively-selected SNVs in lncRNAs ^2,29,30^. For example, *NEAT1* lncRNA, which is a structural component of subnuclear paraspeckle bodies, has been noted for its high mutation rate across a variety of cancers ^29,31,32^. This raises the possibility that a subset of cancer-lncRNAs may also act as “driver-lncRNAs”, where SNVs promote cell fitness by altering lncRNA activity. However, most studies have argued that mutations in *NEAT1* and other lncRNAs arise from phenotypically-neutral passenger effects ^2,29^. To date, the fitness effects of lncRNA SNVs have not been investigated experimentally.

In the present study, we investigate the existence of driver-lncRNAs. We develop an enhanced lncRNA driver discovery pipeline, and use it comprehensively map candidate driver-lncRNAs across the largest cohort to date of somatic SNVs from both primary and metastatic tumours. We evaluate the clinical and genomic properties of these candidates. Finally, we employ a range of functional and mechanistic assays to gather the first experimental evidence for fitness-altering driver mutations acting through lncRNAs.

## Results

### Integrative driver lncRNA discovery with ExInAtor2

Driver genes can be identified by signals of positive selection acting on their somatic mutations. The two principal signals are *mutational burden* (MB), an elevated mutation rate, and *functional impact* (FI), the degree to which mutations are predicted to alter encoded function. Both signals must be compared to an appropriate background, representing mutations under neutral selection.

To search for lncRNAs with evidence of driver activity, we developed *ExInAtor2*, a driver-discovery pipeline with enhanced sensitivity due to two key innovations: integration of both MB and FI signals, and empirical background estimation (see Methods) (Figure 1a, Supplementary Figure 1a, b). For MB, local background rates are estimated, controlling for covariates of mutational signatures and large-scale effects such as replication timing, which otherwise can confound driver gene discovery ^33^. For FI, we adopted functionality scores from the *Combined Annotation Dependent Depletion* (CADD) system, due to its widespread use and compatibility with a range of gene biotypes ^34^. Importantly, *ExInAtor2* remains agnostic to the biotype of genes / functional elements, allowing independent benchmarking with established protein-coding gene data.

**Figure 1.**
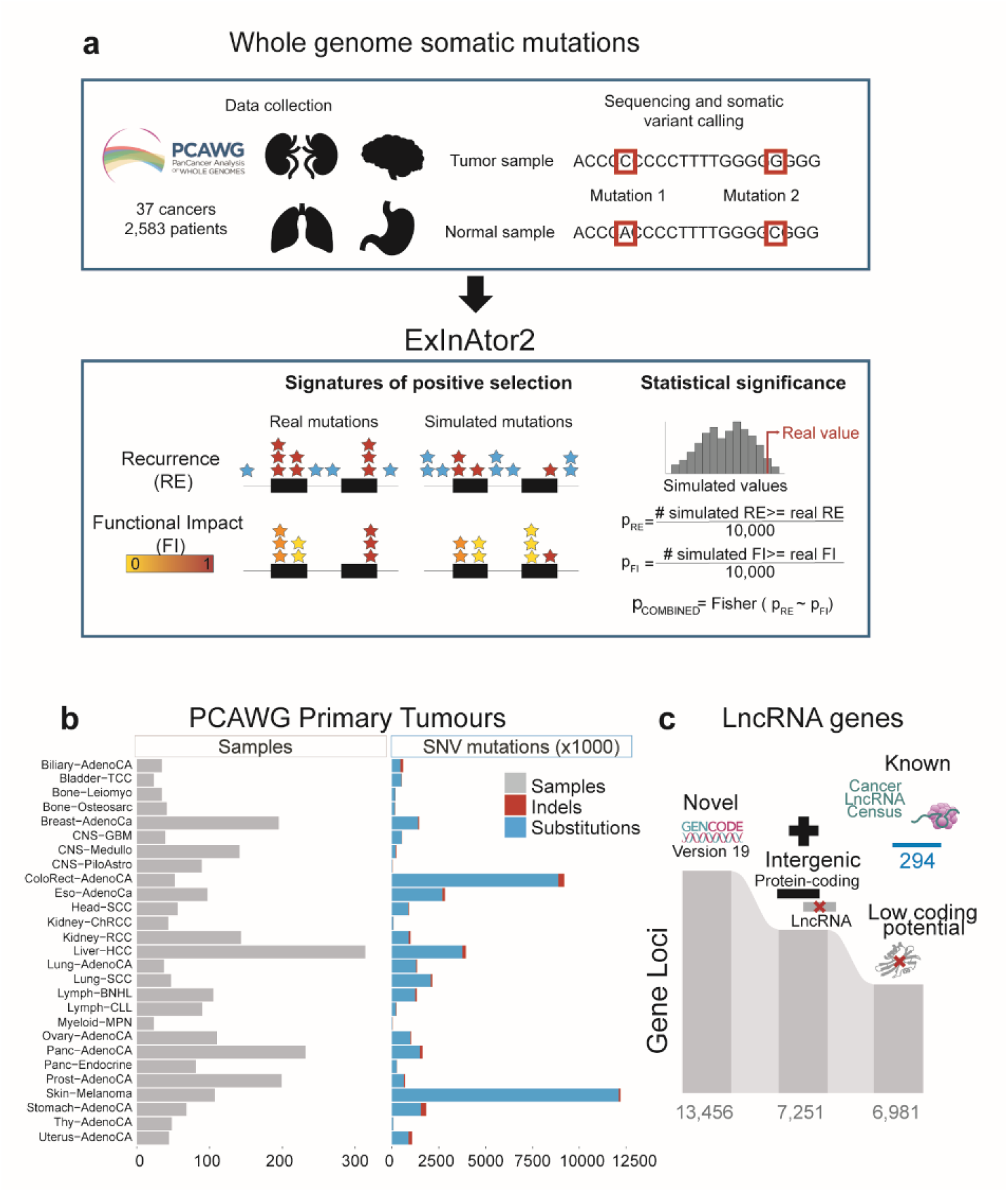
Driver lncRNA discovery with ExInAtor2. **a)** ExInAtor2 accepts input in the form of maps of single nucleotide variants (SNVs) from cohorts of tumour genomes. Two signatures of positive selection are evaluated and compared to simulated local background distributions, to evaluate statistical significance. The two significance estimates are combined using Fisher’s method. **b)** Summary of the primary tumour datasets used here, obtained from Pancancer Analysis of Whole Genomes (PCAWG) project. **c)** A filtered lncRNA gene annotation was prepared, and combined with a set of curated cancer lncRNAs from the Cancer LncRNA Census ^23^.

### Accurate discovery of known and novel driver genes

We began by benchmarking ExInAtor2 using the maps of somatic single nucleotide variants (SNVs) from tumour genomes sequenced by the recent PanCancer Analysis of Whole Genomes (PCAWG) project ^1^, comprising altogether 45,704,055 SNVs from 2,583 donors (Figure 1b, Methods). As it was generated from whole-genome sequencing (WGS), this dataset makes it possible to search for driver genes amongst both non-protein-coding genes (including lncRNAs) and better-characterised protein-coding genes.

To maximise sensitivity and specificity, we prepared a carefully-filtered annotation of lncRNAs. Beginning with high-quality curations from Gencode ^35^, we isolated intergenic lncRNAs lacking evidence for protein-coding capacity. To the resulting set of 6981 genes (Figure 1c), we added the set of 294 confident, literature-curated lncRNAs from Cancer LncRNA Census 2 dataset ^23^, for a total set of 7275 genes.

We compared the performance of ExlnAtor2 to ten leading driver discovery methods and PCAWG’s consensus measure, which integrates and outperforms these individual methods (Figure 2a) ^32^. Performance was benchmarked on curated sets of protein-coding and lncRNA cancer genes (Figure 2b). Judged by correct identification of cancer lncRNAs at a false discovery rate (FDR) cutoff of <0.1, ExInAtor2 displayed the best overall accuracy in terms of F1 measure (Figure 2c, d). Quantile-quantile (QQ) analysis of resulting *p*-values (P) displayed no obvious inflation or deflation and has amongst the lowest Mean Log Fold Change (MLFC) values (Figure 2e), together supporting ExInAtor2’s low and controlled FDR.

**Figure 2.**
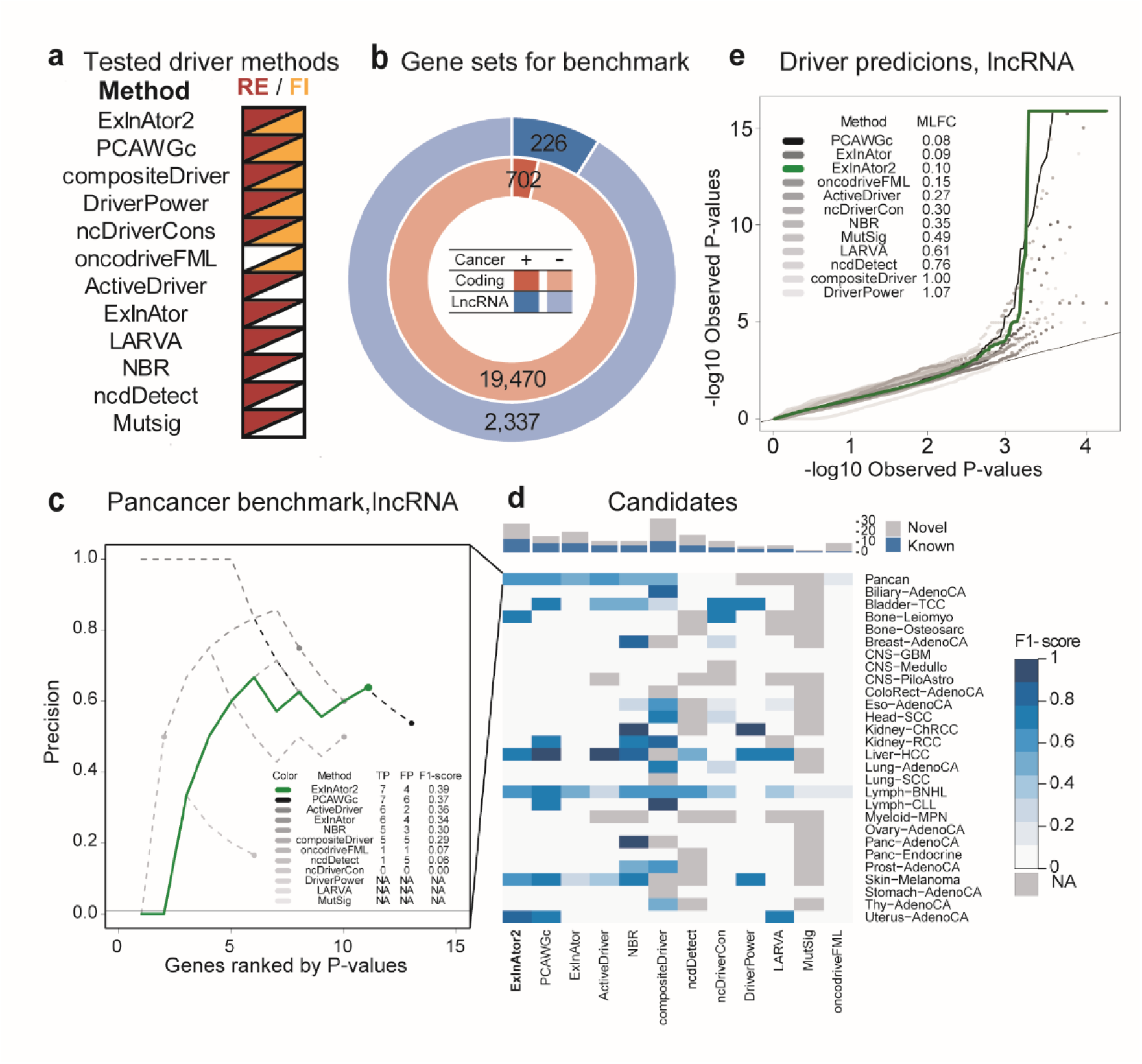
ExlnAtor2 accurately identifies driver genes. **a)** The list of driver discovery methods to which ExInAtor2 was compared. The signatures of positive selection employed by each method are indicated to the right. PCAWGc indicates the combined driver prediction method from Pan-Cancer Analysis of Whole Genomes (PCAWG), which integrates all ten methods. **b)** Benchmark gene sets. LncRNAs (blue) were divided in positives and negatives according to their presence or not in the Cancer LncRNA Census ^23^, respectively, and similarly for protein-coding genes in the Cancer Gene Census ^36^. **c)** Comparing performance in terms of precision in identifying true positive known cancer lncRNAs from the CLC dataset, using PCAWG Pancancer cohort. *x*-axis: genes sorted by increasing *p*-value. *y*-axis: precision, being the percentage of true positives amongst cumulative set of candidates at increasing *p*-value cutoffs. Horizontal black line shows the baseline, being the percentage of positives in the whole list of tested genes. Coloured dots represent the precision at cutoff of *q* ≤ 0.1. Inset: Performance statistics for cutoff of *q* ≤ 0.1. **d)** Driver prediction performance for all methods in all PCAWG cohorts. Cells show the F1-score of each driver method (*x*-axis) in each cohort (*y*-axis). Grey cells correspond to cohorts where the method was not run. The bar plot on the top indicates the total, non-redundant number of True Positives (TP) and False Positives (FP) calls by each method. Driver methods are sorted from left to right according to the F1-score of unique candidates. **e)** Evaluation of *p*-value distributions for driver lncRNA predictions. Quantile-quantile plot (QQ-plot) shows the distribution of observed vs expected –log10 *p*-values for each method run on the PCAWG Pancancer cohort. The Mean Log-Fold Change (MLFC) quantifies the difference between observed and expected values (Methods).

ExInAtor2 is biotype-agnostic, and protein-coding driver datasets are highly refined (Figure 2b). To further examine its performance, we evaluated sensitivity for known protein-coding drivers from the benchmark Cancer Gene Census ^36^. Again, ExInAtor2 displayed competitive performance, characterised by low false positive predictions (Supplementary Figure 2a-c).

To test ExInAtor2’s FDR estimation, we repeated the lncRNA analysis on a set of carefully-randomised pancancer SNVs (see Methods). Reassuringly, no hits were discovered and QQ plots displayed neutral behaviour (MLFC 0.08) (Supplementary Figure 2d). Analysing at the level of individual cohorts, ExInAtor2 predicted 3 / 40 lncRNA-cohort associations in the simulated / real datasets, respectively. This corresponds to an empirical FDR rate of 0.075, consistent with the nominal FDR cutoff of 0.1.

We conclude that ExInAtor2 identifies known driver genes with a low and controlled false discovery rate.

### The landscape of driver lncRNA in primary human tumours

We next set out to create a genome-wide panorama of mutated lncRNAs across human primary cancers. Tumours from PCAWG were grouped into a total of 37 cohorts, ranging in size from two tumours (Cervix-AdenoCa, Lymph-NOS and Myeloid-MDS tumour types) to 314 (Liver-HCC tumour type), in addition to the entire pancancer set (Figure 3a).

**Figure 3.**
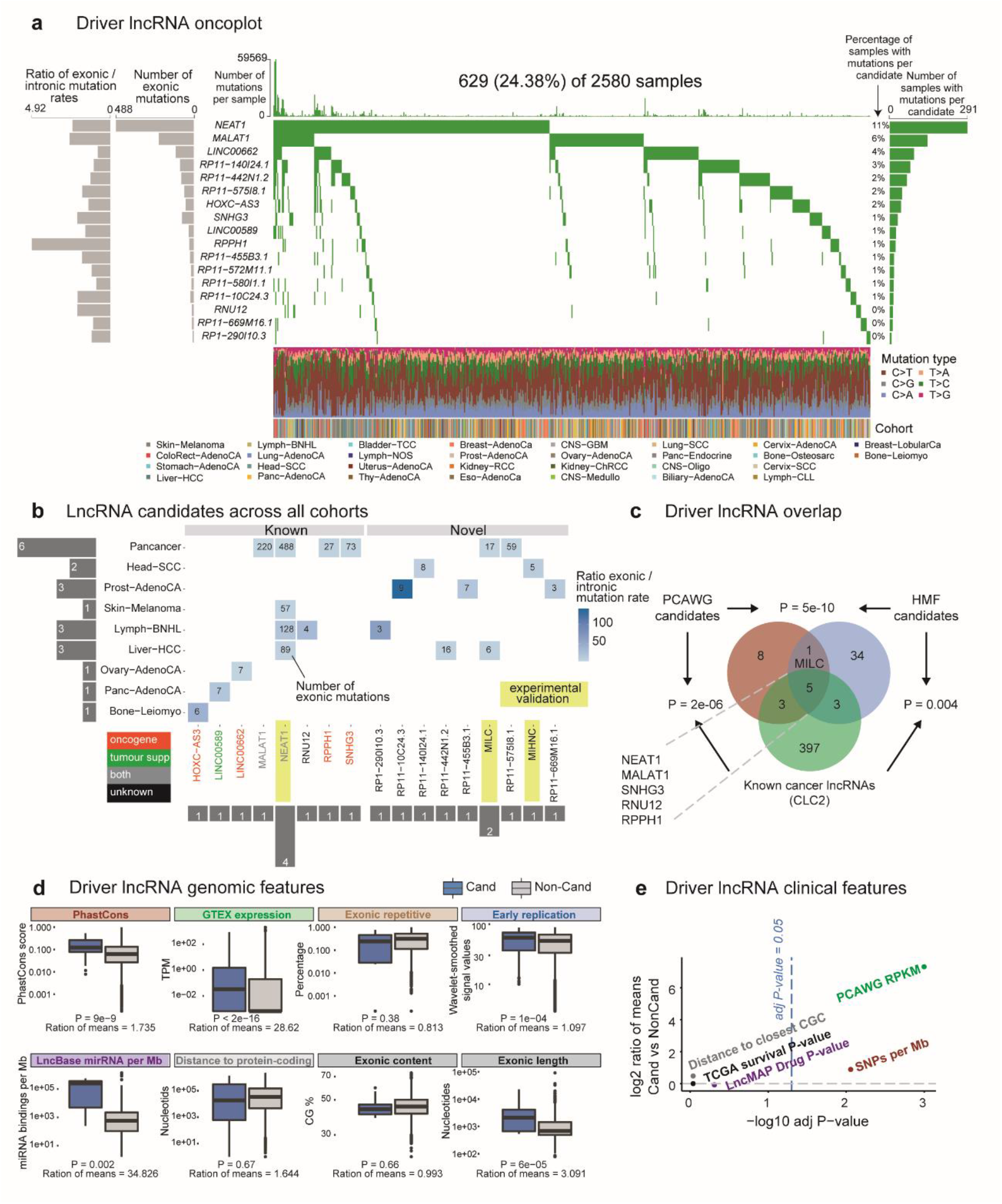
The landscape of known and novel driver lncRNAs in primary tumors. **a)** “Oncoplot” overview of driver lncRNA analysis in PCAWG primary tumours. Rows: 17 candidate driver lncRNAs at cutoff of q ≤ 0.1. Columns: 2580 tumours. **b)** LncRNA candidates across all cohorts. Rows: Cohorts where hits were identified. Columns: 17 candidate driver lncRNAs. “Known” lncRNAs are part of the literature-curated Cancer LncRNA Census (CLC2) dataset ^23^. Functional labels (oncogene / tumour suppressor / both) were also obtained from the same source. **c)** Intersection of candidate driver lncRNAs identified in PCAWG primary tumours, Hartwig Medical Foundation (HMF) metastatic tumours and the CLC2 set. Statistical significance was estimated by Fisher’s exact test. **d)** Genomic features of driver lncRNAs. Each plot displays the values of indicated features for 17 candidate driver lncRNAs (blue) and all remaining tested lncRNAs (non-candidates, grey). Significance was calculated using Wilcoxon test. For each comparison, the ratio of means was calculated as (mean of candidate values / mean of non-candidate values). See Methods for more details. **e)** Clinical features of driver lncRNAs. Each point represents the indicated feature. *y-*axis: log2-transformed ratio of the mean candidate value and mean non-candidate value. *x*-axis: The statistical significance of candidate vs non-candidate values, as estimated by Wilcoxon test and corrected for multiple testing. See Methods for more details.

After removing likely false positive associations using the same stringent criteria as PCAWG ^1^, ExInAtor2 revealed altogether 21 unique cancer-lncRNA associations, involving 17 lncRNAs (Figure 3b) – henceforth considered putative “driver lncRNAs”. Of these, nine are annotated lncRNAs that have not previously been linked to cancer, denoted “novel”. The remaining “known” candidates are identified in the literature-curated Cancer LncRNA Census 2 dataset ^23^. Known lncRNAs tend to be hits in more individual cohorts than novel lncRNAs, with cases like *NEAT1* being detected in four cohorts (Figure 3b). While most driver lncRNAs display exonic mutation rates ∼50-fold greater than background (coloured cells, Figure 3b), the number of mutations in such genes is diverse between cohorts, being Pancancer, Lymph-CLL and Skin-Melanoma the biggest contributors of mutations.

Supporting the accuracy of these predictions, the set of driver lncRNAs is highly enriched for known cancer lncRNAs ^23^ (8/17 or 48%, Fisher test P=2e-6) (Figure 3c). Driver lncRNAs are also significantly enriched in three other independent literature-curated databases (Supplementary Figure 3a).

### Driver lncRNAs carry features of functionality and clinical relevance

To further evaluate the quality of driver lncRNA predictions, we tested their association with genomic and clinical features expected of *bona fide* cancer genes. LncRNA catalogues are likely to contain a mixture of both functional and non-functional genes. The former group is characterised by purifying evolutionary selection and high expression in healthy and diseased tissues ^27^. We found that driver lncRNAs display higher evolutionary sequence conservation and higher steady-state levels in healthy organs (Figure 3d). Their sequence also contains more microRNA binding sites, suggesting integration with post-transcriptional regulatory networks.

In contrast, we could find no evidence that driver lncRNAs are enriched for genomic covariates and features arising from artefactual results. They have earlier replication timing (whereas later replication is associated with greater mutation) ^37^, less exonic repetitive sequence (ruling out mappability biases), and similar exonic GC content (ruling out sequencing bias) compared to tested non-candidates (Figure 3d). However, driver lncRNAs tend to have longer spliced length, likely reflecting greater statistical power for longer genes that affects all driver methods ^29^.

Driver lncRNAs also have clinical features of cancer genes (Figure 3e). They are on average 158-fold higher expressed in tumours compared to normal tissues (133 vs 0.84 FPKM) (Figure 3e, PCAWG RPKM), 2.15-fold enriched for germline cancer-associated small nucleotide polymorphism (SNP) in their gene body (4.7% vs 2.5%) (Figure 3e, SNPs per MB), and enriched in orthologues of driver lncRNAs carrying common insertion sites (CIS), discovered by transposon insertional mutagenesis (TIM) screens in mouse IM screens identify (17.6 vs 1.6%) (Supplementary Figure 3a) ^23^. Finally, driver lncRNAs significantly overlap growth-promoting hits discovered by CRISPR functional screens (11.8 vs 1.3%) (Supplementary Figure 3a). In conclusion, driver lncRNA display evidence for functionality across a wide range of functional and clinical features, strongly suggesting that they are enriched for *bona fide* cancer driver genes.

### The landscape of lncRNA drivers in metastatic tumours

We further extended the driver lncRNA landscape to metastatic tumours, using 3,527 genomes from 31 cohorts sequenced by the Hartwig Medical Foundation (Supplementary Figure 3 b-d) ^38^. Performing a similar analysis as above, we identified 43 driver lncRNAs in a total of 53 lncRNA-tumour combinations (Supplementary Figure 3b). Eight predicted drivers are known cancer lncRNAs, significantly higher than random expectation (P=0.004) (Figure 3c). Further adding confidence to these findings is the significant overlap of driver lncRNAs identified in the metastatic and primary tumour cohorts (Figure 3c).

### Driver mutations identify oncogenic lncRNAs with therapeutic potential

We wished to evaluate the therapeutic and functional relevance of novel lncRNAs identified by driver analysis. ENSG00000241219 (RP11-572M11.1), herein named *MILC* (Mutated in Liver Cancer) displayed elevated mutation rates in Hepatocellular Carcinoma (HCC) tumours (Figure 4a) and has been detected as driver in both the PCWG and HFM datasets. It has, to our knowledge, never previously been implicated in cancer. According to the latest Gencode version 38, its single annotated isoform comprises three exons, and displays low expression in normal tissues (Supplementary Figure 4a). We could detect *MILC* in two HCC cell lines, HuH7 and SNU-475 (Figure 4c and Supplementary Figure 4c). To perturb *MILC* expression, we designed two different antisense oligonucleotides (ASOs) that reduced steady-state levels by >50% in both cell lines (Figure 4b,c and Supplementary Figure 4c). We evaluated the role of *MILC* in HCC cell proliferation, by measuring changes in growth rates following ASO transfection. The significant decrease in growth resulting from both ASOs in both cell backgrounds points to the importance of *MILC* in cell fitness (Figure 4d and Supplementary Figure 4d).

**Figure 4.**
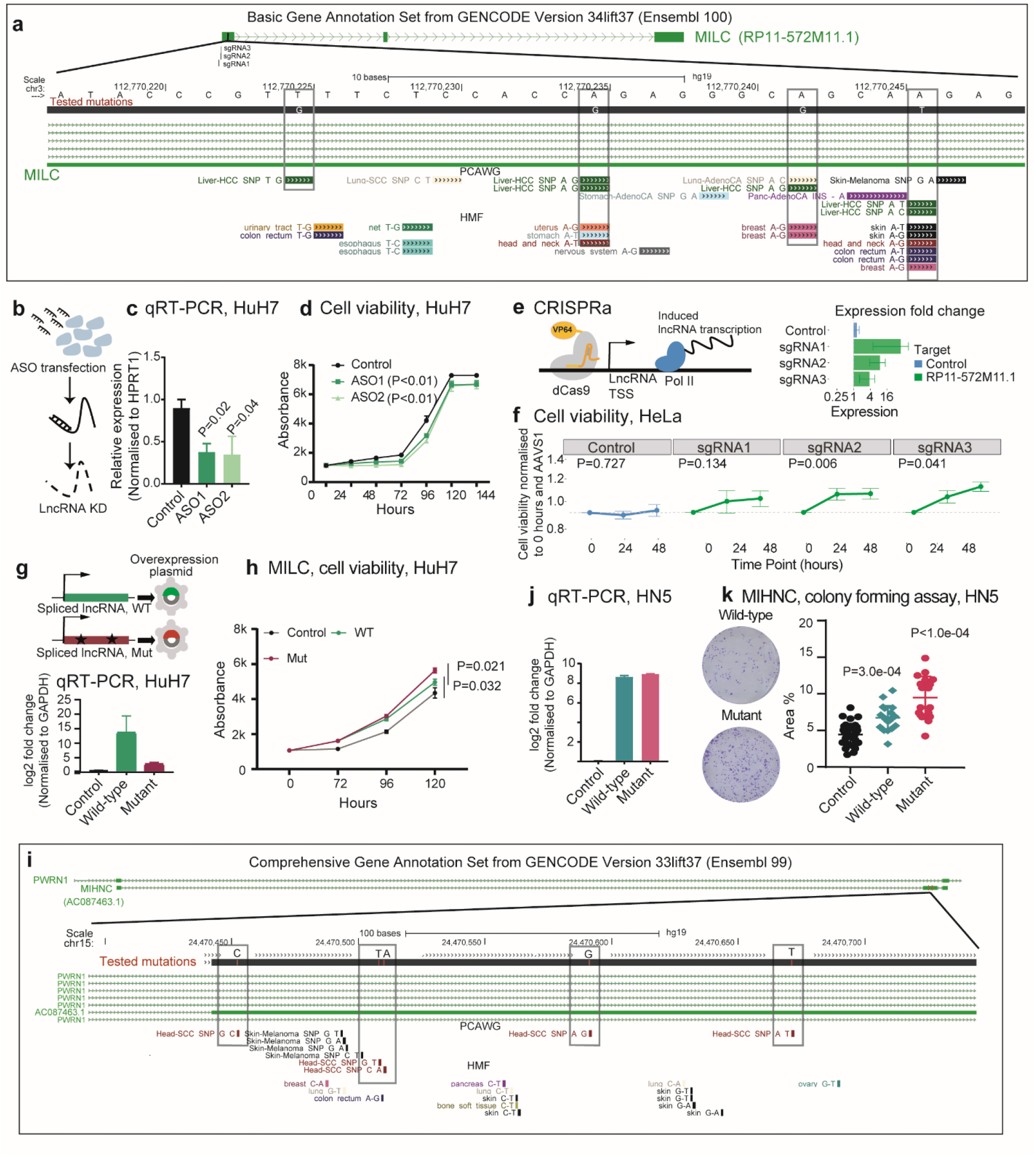
Mutations in *MILC* and *MIHNC* enhance cell fitness. **a)** The genomic locus of hepatocellular carcinoma (HCC) candidate driver lncRNA *MILC*. Also shown are SNVs from PCAWG and Hartwig (HMF). The SNVs included in the mutated version of the plasmids are indicated in the grey boxes. **b)** Antisense oligonucleotides (ASOs) were transfected into cells to knock down expression of target lncRNAs. **c)** Reverse transcription quantitative polymerase chain reaction (qRT-PCR) measurement of RNA levels in HuH HCC cells after transfection of control ASO, or two different ASOs targeting *MILC*. Statistical significance was estimated using one-sided Student’s *t*-test with n=3 independent replicates. **d)** Populations of ASO-transfected cells were measured at indicated time points. Each measurement represents n=3 independent replicates. **e)** Overview and performance of CRISPR-activation (CRISPRa) targeting *MILC*. On the right, qRT-PCR measurements of RNA levels with indicated sgRNAs in HeLa cells. Values were normalised to the housekeeping gene HPRT1 and to a control sgRNA targeting the AAVS1 locus. Values represent n=3 independent replicates. **f)** The effect of CRISPRa on HeLa cells’ viability, as measured by Cell Titre Glo reagent. Values represent n=6 independent replicates, and statistical significance was estimated by comparison to the Control sgRNA by paired *t*-test at the 48 hrs timepoint. **g)** Plasmids expressing spliced *MILC* sequence, in wild-type (WT) or mutated (Mut) form were transfected into HuH cells. The steady state levels of RNA were measured by qRT-PCR and normalised to cells transfected with similar EGFP-expressing plasmid. Values represent n=3 independent replicates, each one with 6 technical replicates. **h)** Populations of plasmid-transfected cells were measured at indicated timepoints. Statistical significance was estimated by one-sided Student’s *t*-test based on n=3 independent replicates. **i)** The genomic locus of head and neck cancer candidate driver lncRNA *MIHNC*. Also shown are SNVs from PCAWG and Hartwig. The SNVs included in the mutated version of the plasmids are indicated in the grey boxes. **j)** Plasmids expressing spliced *MIHNC* sequence, in wild-type (WT) or mutated (Mut) form were transfected into HN5 cells. The steady state levels of RNA were measured by qRT-PCR and normalised to cells transfected with similar EGFP-expressing plasmid. Values represent n=3 independent replicates. **k)** Results of colony formation assay in HN5 cells. Values indicate the percent of well area covered. Statistical significance was estimated using One-way ANOVA has been used to determine statistical significance, based on 18 culture wells.

These results prompted us to ask whether *MILC* can also promote cell growth in other cancer types. Thus, we turned to CRISPR-activation, to upregulate the lncRNA from its endogenous locus in HeLa cervical carcinoma cells. Three independent sgRNAs increased gene expression by 4 to ∼20-fold (Figure 4e and Supplementary Figure 4b), of which two significantly and specifically increased cell proliferation (Figure 4f).

Having established that *MILC* promotes cell growth, we next asked whether tumour mutations can enhance this activity, as would be expected for driver mutations. To do so, we designed overexpression plasmids for the wild-type or mutated forms of the transcript (Figure 4g). The mutated form contained four SNVs, some of them recurrently observed in independent tumours from both PCAWG and HFM dataset (Figure 4a). Transfection of wild-type *MILC* boosted cell growth, consistent with ASO results above. More important, the mutated form resulted in a significant additional increase cell proliferation, compared to the wild-type (Figure 4h).

Another lncRNA, *AC087463*.*1*, herein named *MIHNC* (Mutated in Head and Neck Cancer) was identified as a potential driver in the Head and Neck (HN) tumour cohort (Figure 4i). *MIHNC* is transcribed from the same locus as the lncRNA *PWRN1*, previously reported as a tumour suppressor in gastric cancer 44. It is annotated as a single isoform with three exons (Figure 4i), with the mutations falling in the second, unique exon (Figure 4i). A similar strategy as above showed that overexpression of a mutated form carrying 5 SNVs (Figure 4j) increased tumorigenicity in HN cells, as measured by colony-forming potential (Figure 4k).

Together, these results show that driver analysis is capable of identifying novel oncogenic lncRNAs and, critically, their activity is enhanced by tumour mutations.

### Mutations in NEAT1 promote cell fitness and correlate with survival

To gain mechanistic insights into how fitness-enhancing driver mutations may act through lncRNAs, we turned to a relatively well-understood lncRNA, *NEAT1*, for which confident mechanistic and functional data is available. Based on ExInAtor2 analysis, *NEAT1* mutations, spanning the entire gene length, display evidence for positive selection in altogether 4 and 3 cancer cohorts in PCAWG and Hartwig, respectively. PCAWG and others also noted this highly elevated mutation rate in the *NEAT1* gene, although it has been argued that these result from neutral passenger processes, possibly linked to the high expression of the gene^2,31,40^.

*NEAT1* produces short and long isoforms (called NEAT1_1 / NEAT1_2) of 3.7 and 22.7 kb, respectively ^41^, which are completely overlapping at the 5’ of the gene (Figure 5b). NEAT1_1 is a ubiquitous, abundant, polyadenylated and highly conserved transcript ^42^. In contrast, NEAT1_2, responsible for formation of membraneless nuclear paraspeckle structures, is not polyadenylated and expressed under specific conditions or in response to various forms of stress ^43,44^.

**Figure 5.**
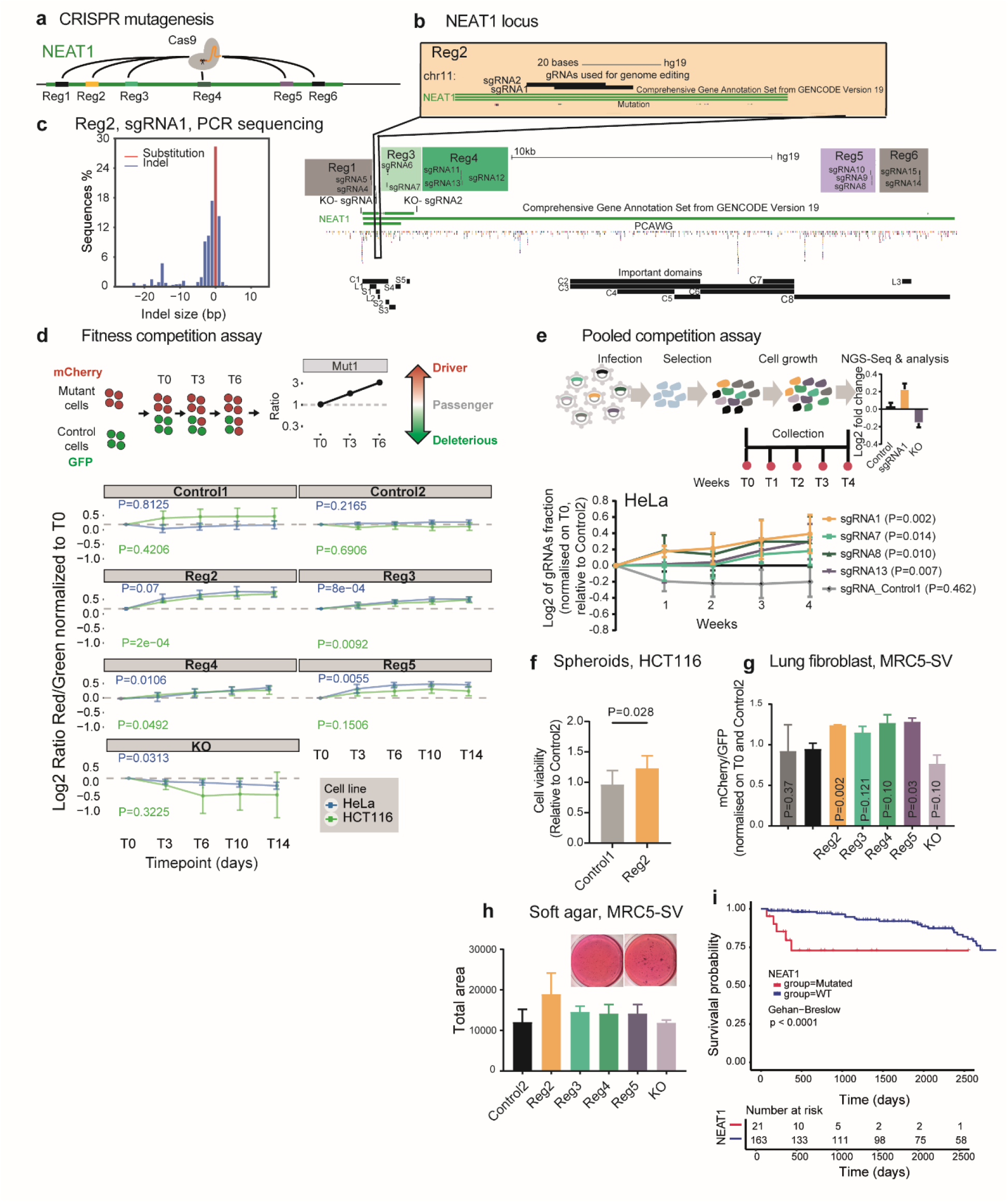
Mutations in *NEAT1* promote cell fitness and correlate with survival. **a)** Overview of the experimental strategy to simulate tumour mutations in the *NEAT1* lncRNA gene by wild-type Cas9 protein. **b)** A detailed map of the six *NEAT1* target regions and 15 sgRNAs. Paired gRNAs used for the deletion of NEAT1_1 are indicated as KO-sgRNA1 and KO-sgRNA2. Previously described functional regions of *NEAT1* are indicated below, according to the publication of Yamazaki and colleagues ^46^. **c)** Analysis of mutations created by Cas9 recruitment. The target region was amplified by PCR and sequenced. The frequency, size and nature of resulting DNA mutations are plotted. **d)** Competition assay to evaluate fitness effects of mutations. Above: Rationale for the assay. Labelled mutated (mCherry, red) and control (GFP, green) cells are mixed in equal proportions at the start of the experiment. At successive timepoints their red/green ratio is measured by flow cytometry, and this value is used to infer fitness effects. Below: Red/green ratios for indicated mutations. “Control1/2” indicate sgRNAs targeting intergenic regions. “KO” indicates paired sgRNAs designed to delete the entire NEAT1_1 region. Separate experiments were performed in HeLa and HCT116 cells. n=4 replicated experiments were performed, and statistical significance was estimated by linear regression model on log2 values. **e)** Upper panel: Setup of mini CRISPR fitness screen. HeLa cells are infected with lentivirus carrying defined mixtures of sgRNAs. The sgRNA sequences are amplified and sequenced at defined timepoints. Changes in abundance reflect effects on cell fitness. Lower panel: Abundances of displayed sgRNAs, normalised to the Control 2 negative control. n=4 independent experiments were performed, and statistical significance was estimated by linear regression model. **f)** HCT116 cells were cultured as spheroids and their population measured. n=4 replicated experiments were performed, and statistical significance was estimated using Student’s one-sided *t*-test. **g)** As for Panel D, but with non-transformed MRC5 lung fibroblast cells at timepoint Day 14. Statistical significance was estimated by one-sided Student’s t-test based on n=3 independent replicates. **h)** MRC5 cells were seeded in soft agar, and the area of colonies at 3 weeks were calculated. The mean of n=2 replicated experiments are shown. **i)** The survival time of 184 lymphoid cancer patients from PCAWG is displayed. Patients were stratified according to whether they have ≥1 SNVs in the *NEAT1* gene.

We sought to test whether indels in *NEAT1* can act as drivers. We hypothesised that tumour indels could be simulated wild-type Cas9 protein, which is known to cause similar mutations when double strand breaks are resolved by error-prone DNA repair pathways ^15,45^. We selected six regions of *NEAT1*, based on high mutation density, evolutionary conservation and known functions ^46^, hereafter called Reg1, Reg2, etc.., and targeted them with altogether 15 sgRNAs (Figure 5a). To control for the non-specific fitness effects of double strand breaks (DSBs) ^47,48^, we also created two neutral control sgRNAs targeting *AAVS1* locus, and a positive-control paired sgRNA (pgRNA) to delete the entire NEAT1_1 region (Figure 5b and Supplementary Figure 5a). Sequencing of treated cells’ gDNA revealed narrowly-focussed substitutions and indels at target regions, similar to that observed in real tumours (Figure 5c and Supplementary Figure 5b).

To quantify mutations’ effects on cell fitness, we established a competition assay between mutated mCherry-labelled cells and control GFP-labelled cells (Figure 5d and Supplementary Figure 5c) ^15^. As expected, deletion of entire NEAT1_1 in HeLa cells led to reduced growth (KO), while control sgRNAs did not (Figure 5d). Notably, HeLa cells carrying *NEAT1* mutations in defined regions displayed increased fitness: two at the 5’ of the gene (Reg2 and Reg3), one internally near the alternative polyadenylation site (Reg4) and one at the 3’ end (Reg5) (blue line, Figure 5d and Supplementary Figure 5c). These findings were supported in 3/4 cases in HCT116 colorectal carcinoma cells (green line, Figure 5d and Supplementary Figure 5c).

To corroborate these findings, we repeated fitness assays in the more complex pooled competition assay. Here, the evolution of defined mixtures of mutant cells is quantified by amplicon sequencing of sgRNA barcodes. Consistent with previous results, cells carrying *NEAT1* mutations outcompeted control cells over time (Figure 5e).

These results were obtained from monolayer cells, whose relevance to real tumours is disputed. Thus, we performed additional experiments in 3-dimensional spheroids grown from mutated HCT116 cells, and observed again that Reg2 mutations led to increased growth (Figure 5f).

The experiments thus far were performed in transformed cancer cells. To investigate whether *NEAT1* mutations also enhance fitness in a non-transformed background, we performed similar experiments in MRC5 immortalised foetal lung fibroblasts. Again, *NEAT1* mutations were observed to increase fitness, in terms of cell growth (Figure 5g) and, at least for Reg2, in terms of anchorage-independent growth (Figure 5h).

We sought independent evidence for the importance of *NEAT1* mutations in real-life cancer progression. Using patient survival data from the PCAWG cohort, we asked whether presence of a *NEAT1* mutation correlates with shorter survival. Indeed, in lymphoid cancer patients, *NEAT1* mutations correlate with significantly worse prognosis (Figure 5i). This effect remains even after accounting for differences in total mutation rates using the Cox proportional hazards model (P=0.02).

In summary, *NEAT1* tumour mutations consistently increase cell fitness *in vitro* independent of genetic background, and are associated with poor prognosis in lymphoid cancer patients.

### Mutations alter the NEAT1 protein interactome and increase paraspeckle formation

*NEAT1* is a necessary component of subnuclear paraspeckles 48,54,55, which assemble when specific architectural proteins bind to nascent NEAT1_2 transcripts ^51^. Paraspeckles are nuclear condensates containing diverse gene regulatory proteins ^43^. They are often observed in cancer cells, ^52^, and are associated with poor prognosis ^53^. Thus, we hypothesised that *NEAT1* mutations might affect cell fitness via alterations in paraspeckle number or structure.

We first evaluated changes in *NEAT1* expression and isoform usage in response to mutations. Mutations caused no statistically-significant change in NEAT1_1 expression, while deletion of NEAT1_1 reduced steady-state levels, as expected (Figure 6a). Interestingly, the only mutation to significantly increase NEAT1_2 levels was in Region 4 (Figure 6b), which is consistent with the fact that it contains the alternative polyadenylation site that mediates switching between the short and long isoforms ^54^.

**Figure 6.**
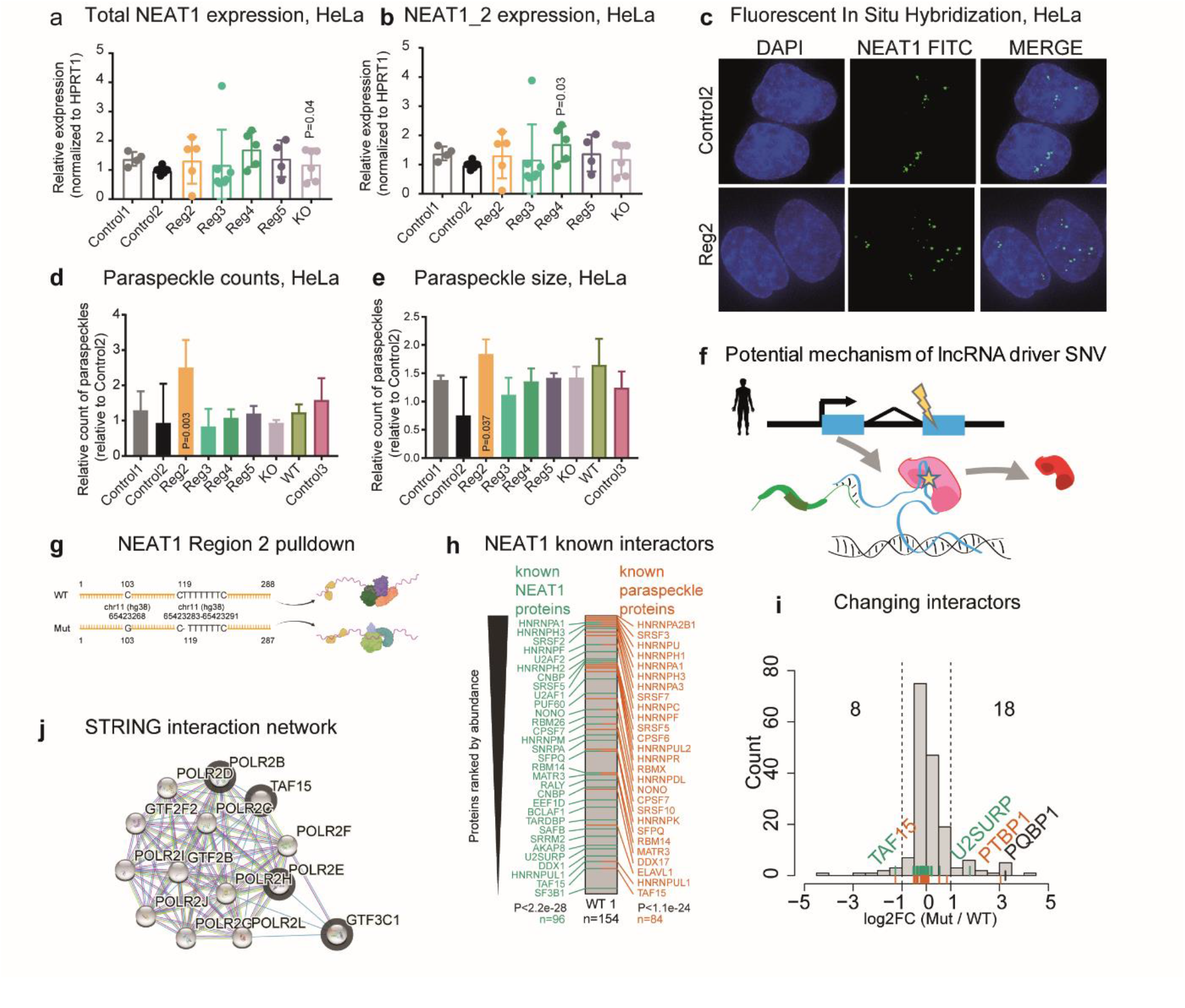
Mutations at the 5’ end of NEAT1 increase paraspeckle formation and alter the protein interactome. **a)** Normalised steady state RNA levels of NEAT1, as estimated using primers for the total NEAT1 region. Statistical significance was estimated using Student’s one-sided *t*-test. P-values ≥0.05 are not shown. **b)** As for Panel A, but using primers for NEAT1_2. **c)** Representative images from fluorescence in situ hybridisation (FISH) visualisation of NEAT1 in HeLa cells expressing sgRNAs for Control 2 and *NEAT1* Region 2. **d)** Counts of paraspeckles in HeLa cells treated with indicated sgRNAs, normalised and compared to Control 2 cells. Values were obtained from 80-100 cells per replicate. N=5 biological replicates. Statistical significance was estimated using paired t-test. **e)** As for Panel D, but displaying paraspeckle size. **f)** Schematic representation of the mechanism of action of driver mutations within *NEAT1* sequence. **g)** Sequences of biotinylated probes used for mass-spectrometry analysis of NEAT1-interacting proteins. **h)** Proteins detected by wild-type (WT) *NEAT1* probe, filtered for nuclear proteins only, are ranked by intensity and labelled when intersecting databases of previously-detected NEAT1-interacting proteins (green) and paraspeckle proteins (orange). Statistical significance was calculated by hypergeometric test (to background of all nuclear proteins n=6758). **i)** Histogram shows differential detection of proteins comparing mutated (Mut) and wild-type (WT) probes. Dotted lines indicate log2 fold-change cutoffs of - 1 / +1. **j)** STRING interaction network based on a subset of the proteins lost upon mutation (grey borders) interacting with the RNA polymerase II core complex.

Using fluorescence in situ hybridisation (FISH) with NEAT1_2 probes, we next asked whether mutations impact on paraspeckle number or structure (Figure 6c). Despite changes in isoform expression noted above, mutations in Region 4 resulted in no change in the number or size of paraspeckles, in line with previous findings ^46^ (Figure 6d,e). However, mutations in Region 2 yielded a significant increase in number and size of paraspeckles (Figure 6c,e).

*NEAT1* is known to function via a diverse cast of protein partners. Region 2 mutations overlap several known protein binding sites, and fall in or near to areas of deep evolutionary conservation of sequence and structure (Supplementary Figure 5d).

To better understand how Region 2 mutations alter *NEAT1* function, and evaluate if mutation could affect the binding of proteins to *NEAT1* (Figure 6f), we compared the protein-interactome of wild-type and mutant RNA by *in vitro* pulldown coupled to mass-spectrometry. We created a 288 nt fragment of NEAT1-Region 2 for wild-type (WT) and mutated sequence, the latter containing two SNVs observed in patient tumours (Figure 6g). We performed RNA pull-down with nuclear lysate from HeLa cells, followed by mass spectrometry. Altogether, 154 interacting nuclear proteins were identified for wild-type sequence. Supporting the usefulness of this approach, interacting proteins highly enriched for both known NEAT1-binders and paraspeckle proteins (see Methods) and include well known examples like NONO ^46,55^ (Figure 6h). Comparing mutant to WT interactomes, we observed widespread changes in *NEAT1* complexes: altogether 8 (4.6%) proteins are lost by mutant RNA, and 18 (10.3%) gained (Figure 6i).

We investigated whether mutations create or destroy known binding motifs of changing proteins, but could find no evidence for this. However, we did note that mutations lead to increased binding of previously-discovered interactors, U2SURP and PTBP1 (Figure 6i). Intriguingly, increased binding was also observed for PQBP1 protein, whose disordered domain has been linked to condensate formation, offering a potential mechanism in facilitating paraspeckle formation ^56^. Conversely, STRING analysis revealed that the proteins lost upon mutation are highly enriched for members of the core RNA Polymerase II complex (strength=2.51, P=0.016; basic list enrichment by STRING, Benjamini-Hochberg corrected) and physically interacting with other proteins of this complex (Figure 6j). In summary, tumour mutations in *NEAT1* give rise to reconfiguration of the protein interactome, creating several potential mechanisms by which paraspeckles formation is promoted in transformed cells.

## Discussion

Understanding which mutations give rise to pathogenic cell fitness, and how they do so, are fundamental goals of cancer genomics. Here we have focussed on a particularly intriguing class of potential driver elements, the lncRNAs, which are known to be both potent cancer genes and highly mutated in tumours, and yet for which no driver mutation has been experimentally validated to date ^2,29,31,57^.

To address this gap, we here developed an improved method, ExInAtor2, to search for driver lncRNAs based on integrated signatures of positive selection. In total, this identified 54 candidate driver lncRNAs across the largest tumour cohort tested to date. The value of these predictions is supported by consistency between independent cohorts, overlap with various cancer lncRNA databases, and from functional screens in mouse. Nevertheless, *in silico* driver analyses suffer from a variety of constraints, from false positives due to localised, non-selected mutational processes, to false negatives due to the limited sample size. Such factors have limited the confidence with which previous studies ^29,30^ could interpret the functional relevance of highly mutated lncRNAs, underlining the importance of experimental results presented here.

The usefulness of novel ExInAtor2 predictions was demonstrated by functional studies on two lncRNAs, *MIHNC* (Head and Neck cancer) and *MILC* (Hepatocellular Carcinoma). Not only are both capable of promoting cancer cell growth in their wild-type form, but interestingly, this activity is enhanced by tumour mutations. These findings provide experimental support for the usefulness of driver analysis in identifying novel cancer lncRNAs.

Among the candidate driver lncRNAs we identified the widely-studied *NEAT1*. Previous tumour sequencing studies have noted the elevated density of SNVs at this locus, but generally attributed them to passenger mutational processes, possibly a consequence of unusually high transcription rate ^2,29,31,57^. Here, we have provided experimental evidence, via naturalistic *in cellulo* mutagenesis with CRISPR-Cas9, that *NEAT1* SNVs reproducibly give rise to increased cell proliferation, in a range of backgrounds including non-transformed cells. The latter raises the intriguing possibility that *NEAT1* SNVs might contribute to early stages of tumorigenesis. Other observations are worthy of mention. Firstly, amongst fitness-altering *NEAT1* SNVs, we only observed those that increase growth, and none that decreased it. Secondly, not all tested regions of NEAT1 could host fitness-altering mutations, and these were clustered at previously-mapped functional elements in mature RNA ^44,46^. Altogether, these findings suggest that tumour SNVs at particular regions of *NEAT1* are phenotypically non-neutral and capable of increasing cell fitness by altering function of encoded RNA. The notion that the *NEAT1* gene represents a vulnerability to tumorigenesis is further supported by our demonstration that patients carrying mutations in the gene have worse prognosis, as well as published transposon insertional mutagenesis screens in mouse ^27^.

The relatively well-understood role of *NEAT1* in assembling ribonucleoprotein phase-separated paraspeckle organelles afforded important insights into SNVs’ molecular mechanisms. Introduction of tumour mutations at the gene’s 5’ end impacted protein binding, including a significant loss of interaction with the RNA Polymerase II complex mediated by known *NEAT1* interactor *TAF15*. Other known protein interactions are potentiated in mutated RNA, suggesting that changes in paraspeckles may be mediated by both gains and losses of protein interactions. The fact that these same mutations gave rise to increased numbers and sizes of paraspeckle structures, suggests a model where SNVs alter the assembly of *NEAT1* ribonucleoprotein complexes, thereby promoting paraspeckle formation and hence cell growth. Future studies will have to address a number of gaps and questions raised here. Firstly, the available of larger tumour cohorts will afford statistical power to discover candidate driver lncRNAs with greater accuracy, while improved statistical models and gene annotations will reduce false positives and false negatives, respectively. While we have provided functional experimental evidence for effects on cell phenotype arising from SNVs, it will be important to replicate this in better models, notably by introducing precise tumour mutations into cellular genomes (eg by recent Prime Editing method)^58,59^, and testing their effects in faithful tumour models, such as mice or tumour organoids ^60,61^. Finally, key mechanistic questions remain to be answered, such as the precise protein partners whose interaction is altered to result in paraspeckle changes.

Phenotype-altering lncRNA mutations could have eventual implications for therapy. We have shown how lncRNA mutations can be prognostic for patient survival, and how driver analysis leads to potential new targets for antisense oligonucleotide (ASO) therapeutics. In future, patients carrying identified driver SNVs in tumour-specific lncRNAs might be treated using personalised cocktails of ASOs, for low-toxicity and effective therapy ^62–64^.

In summary, this work represents the first experimental evidence that fitness-boosting somatic tumour mutations can act via changes in lncRNA function. We have sketched a first mechanistic outline of how this process occurs via altered protein interaction and changes to membraneless organelles, in this case, paraspeckles. Our catalogue of candidate driver lncRNAs across thousands of primary and metastatic tumours provides a foundation for future elucidation of the extent and mechanism of driver lncRNAs.

## Methods

### ExInAtor2 algorithm

ExInAtor2 is composed of two separate modules for detection of positive selection: one for recurrence (RE), comparing the exonic mutation rate to that of the local background; another for functional impact (FI), comparing the estimated functional impact of mutations to background, both estimated in exons.

As an improvement to the first version of ExInAtor ^65^, the RE module compares the number of observed exonic mutations against a distribution of simulated exonic counts (Supplementary Figure 1a), obtained by random repositioning of the variants the between the exonic and background regions while maintaining the same trinucleotide spectrum. Background region is defined for each gene as introns plus 10 kb up and downstream, after removing nucleotides overlapping exons from any other gene. Exonic and background regions can be further filtered to remove any additional “masked” regions defined by the user. In this manuscript, this functionality was used to mask low mappability regions and gap regions obtained from the UCSC Genome Browser (Supplementary File1).

The use of local background and controlling for trinucleotide content is intended to avoid known sources of false positives arising from covariates in mutational processes and mutational signatures, such as replication timing, gene expression, chromatin state, etc ^33^.

A *p*-value is assigned to each gene, being the fraction of simulations with higher or equal number of mutations compared to the observed number (Formula 1).

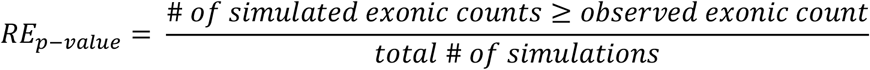

<u>Formula 1: *p*-value calculation for the recurrence (RE) module</u>.

The second FI module compares the mean functional score of the observed exonic mutations to a distribution of simulated values. Simulations are performed by random repositioning of mutations in exonic regions, while maintaining identical trinucleotide content (Supplementary Figure 1b). Similar to the RE model, a *p*-value is obtained by comparing the number of simulations with an exonic mean functional score higher or equal to the observed value (Formula 2). This module work with any base-level scoring method. Given its previous successful use and integrative nature, we selected the Combined Annotation Dependent Depletion (CADD) scoring system ^66^.

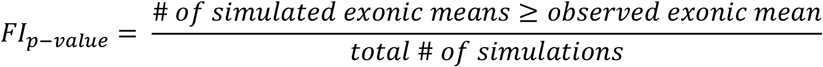

<u>Formula 2: *p*-value calculation for the Functional Impact (FI) module.</u>

In a final step, RE and FI *p*-values are combined using the Fisher method (Formula 3).

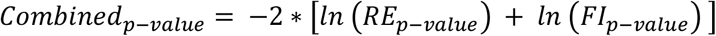

<u>Formula 3: Fisher method for *p*-value integration.</u>

### Tumour somatic mutations

The principal source of mutations were primary tumours from the Pan-Cancer Analysis of Whole Genomes (PCAWG) project ^1^. This dataset was created according to a uniform and strict methodology, including collection of samples, DNA sequencing and somatic variant calling, aggressive filtering to remove potential artefacts and false positive mutations ^1^. For practical reasons, we only considered Single Nucleotide Variants (SNVs) arising from substitutions, insertions and deletions of length 1 bp (indels) (Figure 1b). After this filtering, the PCAWG dataset comprises 37 cancer cohorts, 2,583 samples and 45,703,485 SNVs (Figure 1b). Analyses were performed either on individual cohorts, or on the “Pancancer” union of all cohorts.

### Gene annotation and filtering

We employed a filtered lncRNA gene annotation based upon Gencode annotation. Beginning with Gencode v19 annotation, we discarded lncRNA genes overlapping protein-coding genes, or containing at least one transcript predicted to be protein-coding by CPAT ^67^, with default settings of coding potential >=0.364. To the remaining list of 6981 genes, we added 294 genes from Cancer LncRNA Census (CLC) ^23^, not annotated in Gencode v19. The resulting set of 7275 lncRNA genes were used here unless otherwise specified (Figure 1c; Supplementary File 2).

### ExInAtor2 benchmarking against other driver discovery methods

We collected driver predictions from 10 methods, in addition to the combined predictions generated by the PCAWG driver group (PCAWG combined, PCAWGc) that displayed best overall performance ^2^. We only selected PCAWG methods that were run in both protein-coding and lncRNAs, and for which predictions were available for individual cohorts (Figure 2a).

The original PCAWG publication used carefully filtered annotations for protein-coding and lncRNA genes ^2^. Only coding sequences (CDS) of protein-coding genes were considered, while lncRNAs were strictly filtered by distance to protein coding genes, transcript biotype, gene length, evolutionary conservation and RNA expression. For benchmarking, we ran ExInAtor2 using the same PCAWG annotations.

### Evaluation of *p*-value distributions

Under the assumption that most genes are not cancer drivers and follow the null distribution, the collection of p-values should mimic a uniform distribution with deviation of a small number of genes at very low p-values ^68^. Quantile-quantile plots (QQ-plot) (Figure 2b and Supplementary Figure 3a) display the observed and expected *p*-values in -log10 scale. In order to generate the theoretical distribution for each driver method across all 37 cohorts and the Pancancer set, we ranked the total list of *n* observed p-values from lowest to highest, then for each *i* observed *p*-value we calculated an expected *p*-value according to the uniform distribution (Formula 4).

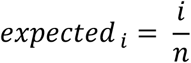

<u>Formula 4: Expected *p*-value calculation. *i* represents the rank of the corresponding observed *p*-value in the total distribution of *n* observed *p*-values, therefore *i* values range from 1 to *n*.</u>

For each driver method, only genes with a reported *p*-value were included in this analysis, i.e., NA cases were discarded. By visual inspection of the QQ-plots, a correct observed distribution of *p*-values should follow a line with 0 as intercept and 1 as slope, where extreme values beyond approximately 2 in the x-axis should deviate above the diagonal line. We used the Mean Log Fold Change (MLFC) (Formula 5) to numerically estimate such deviation and evaluate the performance of driver gene predictions ^68^. The closer to zero the MLFC, the better the statistical modelling of passenger genes following the null distribution ^68^.

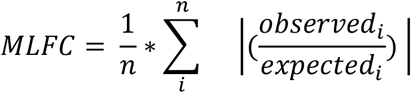

<u>Formula 5: Mean Log Fold Change (MLFC). *n* represents the total number of *p*-values an *i* the lowest *p*-value.</u>

### Gene benchmark sets

We downloaded known driver genes from the Cancer Gene Census ^36^ (CGC) (www.cancer.sanger.ac.uk/census) on 06/02/2019 as a TSV file. We extracted all Gencode *ENSG* identifiers, resulting in a list of 703 genes. For lncRNAs we used the second version of the Cancer LncRNA Census ^23^, which contains 513 Gencode lncRNAs.

### Precision, sensitivity and F1 comparison

CGC and CLC genes were used as ground truth for driver predictions of protein-coding and lncRNAs, respectively. Three metrics were used to compare driver predictions: Precision, the proportion of predictions that are ground truth genes (Formula 6); Sensitivity, the fraction of ground truth genes that are correctly predicted (Formula 7); F1-score, the harmonic mean of precision and sensitivity (Formula 8).

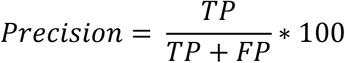

<u>Formula 6: Precision.</u>

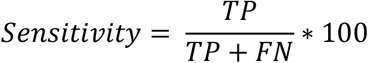

<u>Formula 7: Sensitivity.</u>

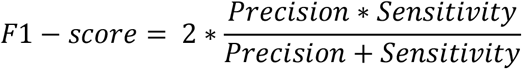

<u>Formula 8: F1-score.</u>

### Simulated mutation datasets

To generate realistic simulated data, each mutation was randomly repositioned to another position with identical trinucleotide signature (ATA > ATA, being the central nucleotide the one mutated) within a window of 50 kb on the same chromosome.

### Generation and comparison of genomic features

Evolutionary conservation: We downloaded base-level PhastCons scores for all 46way and 100way alignments ^69^ from the UCSC Genome Browser ^70^. We calculated the average value across all exons of each gene.

Expression in normal samples: We obtained RNA-seq expression estimates in transcripts per million (TPM) units for 53 tissues from GTEx (https://gtexportal.org/home/datasets). For tissue specificity, we calculated *tau* values as previously described ^71^ (https://github.com/severinEvo/gene_expression/blob/master/tau.R).

Replication timing: We collected replication time data of 16 different cell lines from the UCSC browser ^70^ (http://genome.ucsc.edu/cgi-bin/hgFileUi?db=hg19&g=wgEncodeUwRepliSeq).

miRNA binding: We downloaded both bioinformatically predicted (miTG scores) and experimentally validated miRNA binding to lncRNAs from LncBase ^72^ (http://carolina.imis.athena-innovation.gr/diana_tools/web/index.php?r=lncbasev2%2Findex).

Tumour expression: Expression values in units of FPKM-uq were obtained from PCAWG ^1^.

Drug-expression association: We extracted expression-drug association *p*-values from LncMAP ^73^ (http://bio-bigdata.hrbmu.edu.cn/LncMAP).

Germline cancer small nucleotide polymorphisms (SNPs): We downloaded SNPs from the GWAS Catalogue ^74^ (https://www.ebi.ac.uk/gwas/).

CIS evidence in mice: We downloaded CIS coordinates from CCGD ^75^ (http://ccgd-starrlab.oit.umn.edu/download.php) and mapped them to human hg19 with LiftOver (https://genome.ucsc.edu/cgi-bin/hgLiftOver) from the UCSC browser ^70^. Then, we calculated the number of CIS intersecting each lncRNA divided by the gene length with a custom script using BEDtools ^76^. CIS per Mb values are available in Supplementary File 3.

### Survival analysis

Survival plots were constructed using donor-centric whole genome mutations dataset, overall survival data and tumour histology data from UCSC Xena Hub: https://xenabrowser.net/datapages/?cohort=PCAWG%20(donor%20centric)&removeHub=https://%3A%2F%2Fxena.treehouse.gi.ucsc.edu%3A443. The whole genome mutations file was intersected with comprehensive gene annotation v37 (https://www.gencodegenes.org/human/release_38lift37.html) using BEDtools intersect to isolate donors with mutations in lncRNA of interest. Survival of donors with mutations in lncRNA of interest was then compared against the group of donors without mutations in lncRNA of interest using R packages “survival” (https://cran.r-project.org/web/packages/survival/index.html) and “survminer” (https://cran.r-project.org/web/packages/survminer/index.html)

### NEAT1 structure and element analysis

Elements: The window spanning 300 bp around Mut1a and Mut1b (hg19 chr11:65190589-65190888; hg38 chr11:65423118-65423417) was annotated with the program ezTracks ^77^ using the following datasets as input: (i) structural features: RNA structures conserved in vertebrates (CRS) ^78^, DNA:RNA triplex structures ^79^, R-Loops lifted over to hg38 ^80^; (ii) conservation: phastCons conserved elements in 7, 20, 30 and 100-way multiple alignments ^69^ retrieved from UCSC genome browser ^81^; (iii) high confidence narrow peaks from eCLIP experiments from ENCODE ^82^ (Complete list of accessions is located at Supplementary Table 2).

RBP motif mapping. The 20 bp-padded sequence around Mut1a and Mut1b (hg19 chr11:65190719-65190775) was extracted and then used to generate the sequence of the three distinct alleles WT, only Mut1a and only Mut1b. The three sequences were used as input for de novo RBP motif matching in the web servers RBPmap ^83^ using the option Genome: other and all Human/Mouse motifs) and RBPDB ^84^ (using the default score threshold, 0.8). Outputs were manually parsed and further processed using an in-house Python script.

SNP structural impact analysis. Sequences for the window spanning 300 bp around each mutation target were extracted. Then, only substitutions were kept and encoded according to their relative position and submitted to the MutaRNA web server ^85^, which also reports scores from RNAsnp ^86^.

### Cell culture

HeLa, HEK 293T and HCT116 were a kind gift from Roderic Guigo’s lab (CRG, Barcelona). The MRC5-SV cells were provided by the group of Ronald Dijkmanthe (Institute of Virology and Immunology, University of Bern) and the HN5 tongue squamous cell carcinoma cells by Jeffrey E. Myers (MD Anderson) to Y. Zimmer. All the cell lines were authenticated using Short Tandem Repeat (STR) profiling (Microsynth Cell Line Typing) and tested negative for mycoplasma contamination.

HeLa, HN5 and HEK 293T cell lines were cultured at 37°C in 5% CO2 in Dulbecco’s Modified Eagle’s Medium high-glucose (Sigma) supplemented with: 10% FBS (Gibco), 1% L-Glutamine (ThermoFisher), 100 I.U./mL of Penicillin/Streptomycin (Thermo Fisher).

HCT-116 and MRC5-SV were cultured in McCoy (Sigma) and EMEM (Sigma), respectively, both supplemented with 10% FBS (Gibco), 1% L-Glutamine (ThermoFisher), 100 I.U./mL of Penicillin/Streptomycin (Thermo Fisher). SNU-475 (ATCC) and HuH7 (Cell Line Service) hepatocellular carcinoma cell lines were cultured at 37°C in 5% CO2 in RPMI-1640, GlutaMAX™ (Gibco) supplemented with 10% FBS (Gibco) and 100 I.U./mL of Penicillin/Streptomycin (Thermo Fisher).

### Gene overexpression and knockdown experiments

Both the wild-type and mutated lncRNA spliced sequences were synthesized by Gene Universal Inc, into pcDNA3.1 vector backbone. Control pcDNA3.1 plasmids contained the sequence of enhanced green fluorescent protein (EGFP).

Overexpression in HN5 cells: For each transfection 1.6 ug of plasmid DNA has been incubated for 20 minutes with 4 µl of Lipofectamine 2000 transfection reagent (Invitrogen) in 0.2 ml of OptiMEM media (Gibco) and added to the cells cultured in a 6-well plate. As all plasmids contain G418 resistance gene, cells were cultured in 2.5 mg/ml of G418 (Gibco) 48h after transfection.

Overexpression in HuH7 cells: For each transfection, 100 ng of plasmid DNA were incubated for 20 minutes with 0.15 μl Lipofectamine 3000 and 0.2 μl P3000 transfection reagent (Invitrogen) in 10 μl RPMI-1640, GlutaMAX™ (Gibco) and added on top of 2000 HuH7 cells cultured in a 96-well plate. Transfection efficiency was measured with qPCR after 120h. Knockdown in SNU-475 and HuH7 cells: For the transfections, 10 nM of each ASO were incubated with 0.15 μl Lipofectamine 3000 (Invitrogen) for 20 min in 10 μl RPMI-1640, GlutaMAX™ (Gibco) and added on top of 2000 SNU-475 or HuH7 cells cultured in a 96-well plate. Transfection efficiency was measured with qPCR after 144h.

ASO sequences available in Supplementary File 4.

### Crystal violet staining

Cells were dissociated with 0.05% trypsin-EDTA (Gibco), resuspended in complete media and counted in Neubauer chamber. Subsequently, 1000 cells per well were plated in a 6-well plate, cultured for one week and stained in a 2% Crystal violet (Sigma) solution. The area percentage covered with cells was analysed using ImageJ (%Area). Data analysis was conducted in Graphpad Prism version 8.0.1. One-way ANOVA was used to determine statistical significance, alpha=0.05.

### Proliferation assay – SNU-475 and HuH7

After transfection, the proliferative capacity of SNU-475 and HuH7 was measured every 24h by resazurin assay. Briefly, Resazurin sodium salt (Sigma) was added to each well to reach a final concentration of 3 μM and was incubated at 37°C for 2h. Absorbance was measured with Tecan Spark Plate Reader at 545 nm and 590 nm.

### CRISPR sgRNA design and cloning

CRISPR activation in HeLa cells was performed as described by Sanson and colleagues ^87^. sgRNAs were designed using the GPP sgRNA Designer CRISPRa from the Broad Institute (https://portals.broadinstitute.org/gpp/public/) (Supplementary File 4). For each sgRNA, forward and reverse DNA oligos were synthesized introducing the BsmB1 overhangs. The two oligos were phosphorylated with the Anza™ T4 PNK Kit (Thermofisher) according to the manufacturer instructions in a 10 µl final volume. The phosphorylation/annealing reaction was set up in a thermocycler at 20° C for 15 min, followed by 95°C for 5 min and then ramp down to 25° C at 5° C/min rate. For ligation of annealed oligos into the pXPR_502 backbone (Addgene #96923), the plasmid was first digested and dephosphorylated with FastDigest BsmBI and FastAP (Thermofisher) at 37°C for 2 hrs. Ligation reaction was carried out with the Rapid DNA Ligation Kit (Thermo) according to the manufacturer instructions.

sgRNAs targeting *NEAT1* were designed using the GPP sgRNA Designer CRISPRKo from the Broad Institute (https://portals.broadinstitute.org/gpp/public/) (Supplementary File 4), and cloned into the pDECKO backbone (Addgene #78534) as described above.

### Lentivirus production

For lentivirus production, HEK293T cells (2.5 x10^6) were seeded in poly-L-lysine coated 100 mm culture dishes 24 hrs prior to transfection. Cells were then co-transfected in serum-free medium with 12.5 µg of the plasmid of interest (Lenti dCAS-VP64_Blast plasmid or sgRNA-containing pXPR_502 or pDECKO), 4 µg of the envelope-encoding plasmid pVSVg (Addgene 12260) and 7.5 µg of the packaging plasmid psPAX2 (Addgene 8454) with Lipofectamine 2000 (ThermoFisher) according to the manufacturer instructions. After 4-6 hrs the medium was replaced with complete DMEM. Virus-containing supernatant was collected after 24, 48 and 72 hours post-transfection. The three harvests were pooled and centrifuged at 3000 rpm for 15 min to remove cells and debris. The supernatant was collected, and for every four volumes, one volume of cold PEG-it Virus Precipitation Solution was added. The mix was refrigerated overnight at 4°C and centrifuged at 1500 × g for 30 min at 4°C.The supernatant was discarded, and the sample centrifuged at 1500 × g for 5 min. The lentiviral pellet was suspended in cold, sterile PBS, aliquoted into cryogenic vials and stored at -70°C.

### Lentivirus transduction

#### CRISPRKo

For the generation and transduction of Cas9-expressing cell lines, HeLa, HCT116 and MRC5-SV Cas9 were incubated for 24 hrs with culture medium containing concentrated viral preparation carrying pLentiCas9-T2A-BFP and 8 μg/ml Polybrene. 24 hrs post-infection, antibiotic selection was induced by supplementing the culturing medium with 4 μg/ml blasticidin (Thermofisher) for 5 days. Blasticidin selected cells were subjected to 3 rounds of fluorescence-activated cell sorting (FACS) to isolate high BFP-expressing cells.

#### CRISPRa

For the generation and transduction of dCas9-expressing cell lines, HeLa cells were incubated for 24 hrs with culture medium containing concentrated viral preparation carrying pLenti dCas9-T2A-BFP-VP64 and 8 μg/ml Polybrene. Cells underwent FACS sorting to enrich for high BFP expressing cells.

#### sgRNAs

pLentiCas9-T2A-BFP or dCas9-T2A-BFP-VP64 stable cell line were seeded into 6 well plates at 10^6 cells per well and supplemented with sgRNAs pDECKO or pXPR_502 lentiviral preps, respectively, and spinfected in the presence of polybrene (2 μg/ml) for 95 min at 2000 rpm at 37 °C, followed by medium replacement. 24 hrs post-infection, antibiotic selection was induced by supplementing the culturing medium with 2 μg/ml puromycin (Thermofisher) for at least 3 days.

### RT-qPCR gene expression analysis

HeLa cells were lysed, and total RNA was extracted by using the Quick-RNA™ Miniprep Kit (Zymo Research). For each sample, RNA was retro-transcribed into cDNA by using the GoScript™ Reverse Transcription System (Promega) and the expression of the target gene was assessed through Real-Time PCR with the GoTaq® qPCR Master Mix. To this purpose target-specific mostly intron-spanning primers (Supplementary File 4) were designed by using the online tool Primer 3 version 4.1.0.

### Cell viability assay

After puromycin selection, cells expressing controls and candidates’ guides were collected and seeded in 96-well plates in at least 3 technical replicates for each time point (3000 cells per well). Proliferation assay was performed using the Cell-Titer Glo 2.0 (Promega) reagent according to the manufacturer instructions. Luminescence was measured with the INFINITE 200 PRO series TECAN reader instrument. Time point 0 (T0) reading was performed 4-5 hours after cell seeding.

### 1:1 competition assay

HeLa, HCT116 and MRC5-SV cells were infected with pDECKO lentiviruses expressing fluorescent proteins. Control plasmids containing sgRNAs targeting *AAVS1* expressed GFP protein (pgRNAs-AASV1-GFP+), while the sgRNAs targeting the different regions of *NEAT1* expressed mCherry. After infection, and seven days of puromycin (2 μg/ml) selection, GFP and mCherry cells were mixed 1:1 in a six-well plate (150,000 cells). Cell counts were analysed by LSR II SORP instrument (BD Biosciences) and analysed by FlowCore software.

### Pooled competition assay

Screen: HeLa cells stably expressing sgRNAs targeting *NEAT1* Reg2, Reg3, Reg4, Reg5 and KO, and HeLa cells stably expressing sgRNAs Control1 and Control2 were counted and mixed in the following ratio 10:10:10:10:25:25. At Day 0, 2M cells were collected, while 2M were plated and passaged every 2-3 days. Cells were harvested at 7, 14, 21 and 28 days for gDNA extraction. The experiment was conducted in six biological replicates.

Genomic DNA preparation and sequencing: Genomic DNA (gDNA) was isolated using the Blood & Cell Culture DNA Mini (<5e6 cells) Kits (Qiagen, cat. no. 13323) as per the manufacturer’s instructions. The gDNA concentrations were quantified by Nanodrop. For PCR amplification, 1 μg of gDNA was amplified in a 200 μl reaction using Q5® High-Fidelity 2X Master Mix (NEB #M0491). PCR master mix (100 μl Q5, and 10 μl of Forward universal primer, and 10 μl of a uniquely barcoded P7 primer (both stock at 10 μM concentration). PCR cycling conditions: an initial 30 sec at 98 °C; followed by 10 sec at 98 °C, 30 sec at 68 °C, 20 sec at 72 °C, for 22 cycles; and a final 2 min extension at 72 °C. NGS primers are listed in Supplementary File 4. PCR products were purified with Agencourt AMPure XP SPRI beads according to manufacturer’s instructions (Beckman Coulter, cat. no. A63880). Purified PCR products were quantified using the Qubit™ dsDNA HS Assay Kit (ThermoFisher, cat. no. Q32854). Samples were sequenced on a HiSeq2000 (Illumina) with paired-end 150 bp reads. The raw sequencing reads from individual samples were analysed by using a custom shell script to count the number of reads containing each sgRNA. The sgRNA counts were then normalized over the T0 and Control2.

### Deep sequencing to determine indel spectrum

Genomic DNA was extracted using the Blood & Cell Culture DNA Mini (<5M cells) Kits (Qiagen, cat. no. 13323) as per the manufacturer’s instructions. To prepare samples for Illumina sequencing, a two-step PCR was performed to amplify the different regions of *NEAT1*. For each sample, we performed two separate 100 ul reactions (25 cycles each) with 250 ng of input gDNA using Q5 MASTER MIX (NEB #M0491) and the resulting products were pooled (PCR reaction: 30 sec at 98 °C; followed by 10 sec at 98 °C, 30 sec at 68 °C, 20 sec at 72 °C, for 22 cycles; and a final 2 min extension at 72 °C). PCR amplicons were purified using solid phase reversible immobilization (SPRI) beads, run on a 1.5% agarose gel to verify size and purity, and quantified by Qubit Fluorometric Quantitation (Thermo Fisher Scientific). The resulting DNA was used for reamplification with primers containing Illumina adaptors using the Q5 master Mix. Illumina adaptors and index sequences were added to 100 ng of purified PCR amplicon (PCR reaction: 30 sec at 98 °C; followed by 10 sec at 98 °C, 30 sec at 68 °C, 20 sec at 72 °C, for 8 cycles; and a final 2 min extension at 72 °C).

### RNA-FISH and immunofluorescence

HeLa cells grown on coverslips were fixed using 4% paraformaldehyde and permeabilised by 70% ethanol overnight. For RNA-FISH, Stellaris® FISH Probes, targeting Human *NEAT1* Middle Segment, labelled with FAM dye (1:100, Biosearch Technologies) were used and the procedure was carried out according to the manufacturer’s instructions. Cells nuclei were counterstained with 1:15,000 DAPI (4′,6-diamidino-2-phenylindole) at room temperature and then mounted onto slides by using the VectaShield (Vector Laboratories) mounting media. Fluorescence signals were imaged at 100× (UPLS Apo 100×/1.40) using the DeltaVision Elite Imaging System and Softworx software (GE Healthcare). Images were acquired as Z-stacks, subjected to deconvolution, and projected with maximum intensity. Images were processed using a custom CellProfiler pipeline to determine paraspeckle number and size.

### Soft agar assay

The soft agar colony formation assay was performed as previously described (Borowicz S., et al., 2014). Briefly, the assay was carried out in 6-well plates coated with a bottom layer of 1% noble agar in 2X DMEM (ThermoFisher) supplemented with: sodium bicarbonate, 10% FBS (Gibco), 1% L-Glutamine (ThermoFisher), 100 I.U./ml of Penicillin/Streptomycin (ThermoFisher). Then, 7000 cells were suspended in 2X DMEM and 0.6% noble agar. The suspension mixture was subsequently applied as the top agarose layer. A layer of growth medium was added over the upper layer of agar to prevent desiccation. The plates were incubated at 37 °C in 5% CO2 for 3 weeks until colonies formed. After 20 days the colonies were stained with 200 ml of MTT [(3-(4,5-dimethylthiazol-2-yl)-2,5-diphenyltetrazolium bromide), (5 mg/ml), Sigma] and incubated for 3 hours at 37 °C. Numbers of colonies were counted using the analysis software ImageJ.

### 3D spheroid assay

HCT116 stably expressing Cas9-BFP and sgRNA-mCherry targeting *NEAT1* locus were FACS sorted to enrich the population BFP+/mCherry+. The cells were allowed to grow for 7 days, then detached, counted and seeded onto Corning® 96-well Flat Clear Bottom White (Corning, cat. no. 3610) in 20 μl domes of Matrigel® Matrix GFR, LDEV-free (Corning, cat. no. 356231) and McCoy (Sigma, cat. No. M9309) growth medium (1:1) with a density of 10,000 cells per dome in four technical replicates. Matrigel containing the cells was allowed to solidify for an hour in the incubator at 37 °C before adding 80ul of McCoy growth media on top of the wells. The spheroids were allowed to grow in the incubator at 37°C in a humid atmosphere with 5% CO2. After 4 h the number of viable cells in the 3D cell culture was recorded as time point 0 (T0), CellTiter-Glo® 3D Cell Viability Assay (Promega, cat. no. G9682) was added to the wells, following the manufacturer’s instructions for the reading with the Tecan Infinite® 200 Pro. After one week the measurement was repeated.

### RNA pull-down and Mass Spectrometry

RNA pull-down analysis was performed as previously described (Marín-Béjar O, Huarte M., 2015). Briefly, wild-type and mutant *NEAT1* RNA fragments were transcribed in vitro using HiScribe™ T7 High Yield RNA Synthesis Kit (NEB, #E2040S) and labelled with Biotin using Biotin RNA Labelling Mix (Roche, #11685597910) according to the manufacturers’ instructions. Biotinylated RNA (10 pmol) was denatured for 10 min at 65 °C in RNA Structure Buffer (10 mM tris-HCl, 10 mM MgCl2, and 100 mM NH4C1) and slowly cool down to 4 °C. Nuclear fractions were collected as described previously (Carlevaro-Fita J., et al., 2018) and precleared for 30 min at 4 °C using Streptavidin Mag Sepharose® (Sigma, #GE28-9857-99) and NT2 Buffer [50 mM tris-HCl (pH 7.4), 150 mM NaCl, 1 mM MgCl2, 0.05% NP-40,1 mM DTT, 20 mM EDTA, 400 mM vanadyl-ribonucleoside, RNase inhibitor (0.1 U/µl; Promega), and l× protease inhibitor cocktail (Sigma)]. The precleared nuclear lysates (2 mg) were incubated with purified biotinylated RNA in NT2 buffer along with Yeast tRNA (20 µg/ml; Thermo Fisher Scientific #AM7119) with gentle rotation for 1.5 hours at 4°C. Washed Streptavidin Magnetic Beads were added to each binding reaction and further incubated at 4 °C for 1 h to precipitate the RNA-protein complexes. Beads were washed briefly five times with NT2 Buffer, and the retrieved proteins were then subjected to mass spectrometry analysis, performed by the Proteomics & Mass Spectrometry Core Facility (PMSCF) of the University of Bern, Switzerland, using MaxQuant software for protein identification and quantification.

### Mass Spectrometry Data Processing

Intensity Based Absolute Quantification (iBAQ) and label-free quantitation (LFQ) intensities from the MaxQuant output were used for quantitative within-sample comparisons and fold-enrichment between-sample comparisons respectively. A protein was considered enriched / depleted in a sample condition if its intensity was at least 2-fold greater / lesser than in the reference condition (proteins not detected in one of the conditions are imputed with the lowest value for that sample by MaxQuant). Additionally, the resulting lists of proteins were filtered for nuclear localization ^88^ to exclude potential false positives. To calculate the significance of the overlap with known *NEAT1* binding proteins ^89–91^ and known paraspeckle proteins ^43^ a hypergeometric test was applied to the background of all nuclear proteins (n=6758). STRING was used for interaction analysis (physical subnetwork, minimum interaction score=0.4, max number of direct interactors=10) and GO term enrichment analysis^92^. Visualization of the results was done with R version 4.1.1 and BioRender.com.

## Code availability

The code is accessible at: https://github.com/gold-lab/ExInAtor2.git

## Acknowledgements

The results shown here are based upon data generated by the TCGA, PCAWG and GTEx consortia. We thank Iñigo Martincorena (Sanger Institute) for generously providing certain data analysis scripts. We thank Federico Abascal (Sanger Institute) for generously providing cancer cell fraction data. We thank Adrian Ochsenbein, Carsten Riether, Simon Haefliger, Thomas Marti, Renwang Peng, (Inselspital University Hospital of Bern) for many insightful conversations. We thank Basak Ginsbourger (DBMR) for administrative support, and Willy Hofstetter and Patrick Furer (DBMR) for logistical support. All computation was performed on the Bern Interfaculty Bioinformatics Unit computing cluster maintained by Rémy Bruggmann and Pierre Berthier. This publication and the underlying study have been made possible partly on the basis of the data that Hartwig Medical Foundation has made available. Work in the Johnson laboratory is funded by the Medical Faculty of the University of Bern, the University Hospital of Bern, the Helmut Horten Stiftung, Swiss Cancer Research Foundation (4534-08-2018), Science Foundation Ireland through Future Research Leaders award 18/FRL/6194, and the Swiss National Science Foundation through the National Centre of Competence in Research (NCCR) “RNA & Disease”.

