## Supplementary figures and images for "Tumour mutations in long noncoding RNAs enhance cell fitness"

### Supplementary Figure 1

Supplementary Figure 1

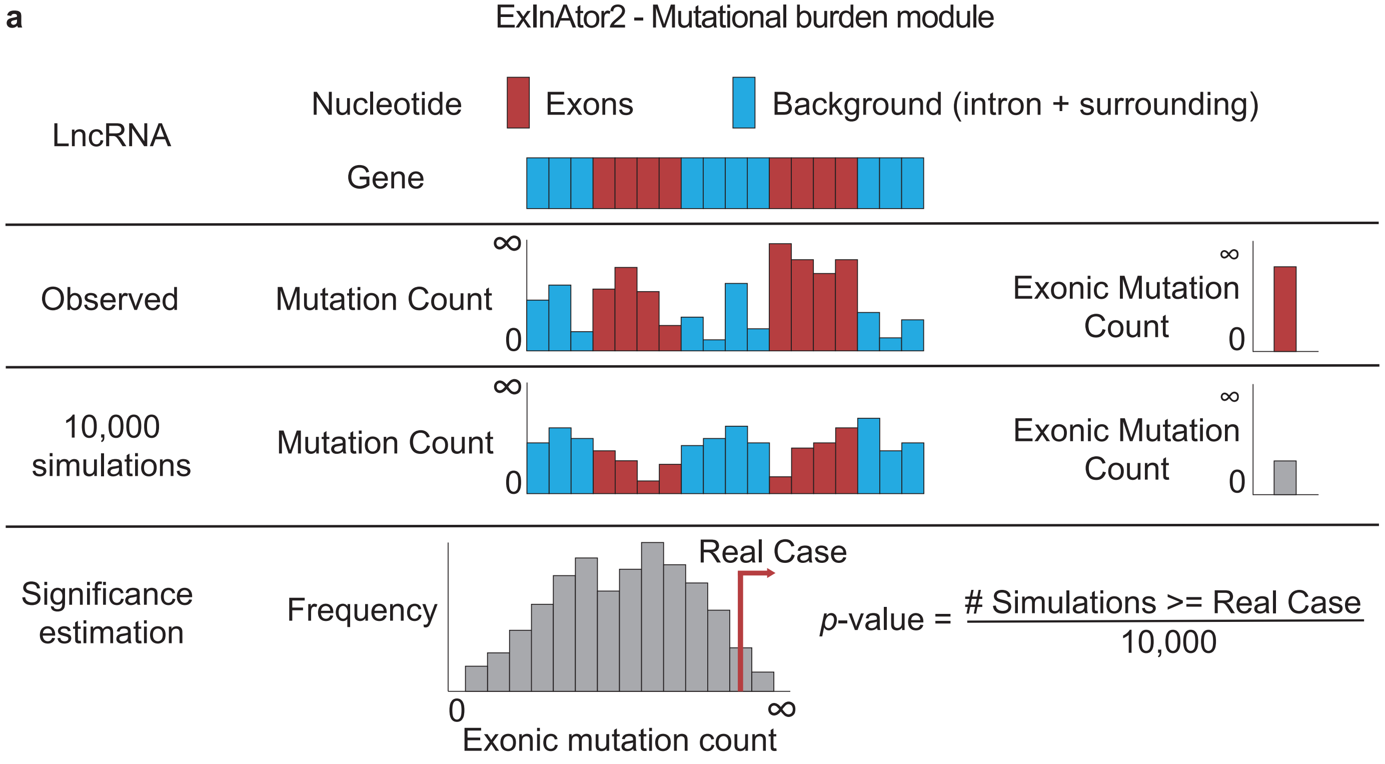


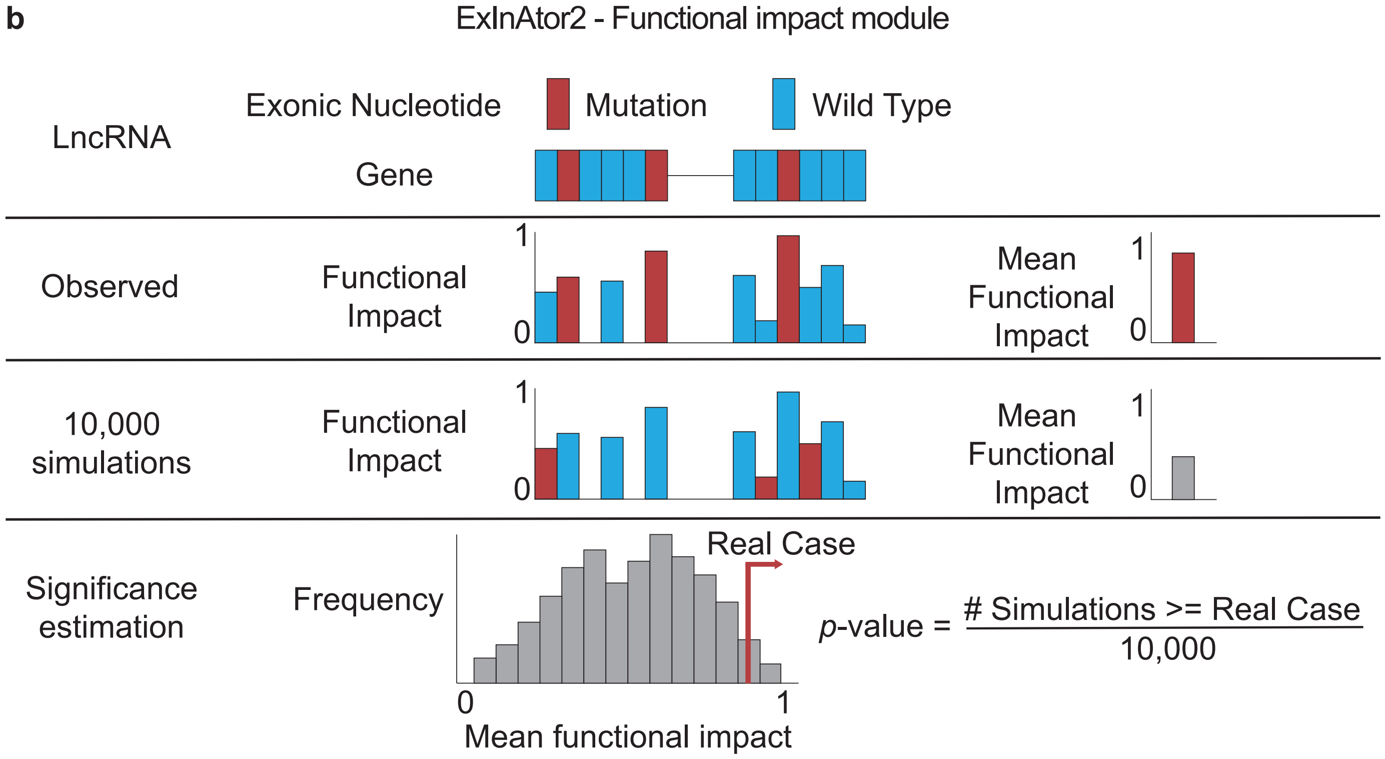

### Supplementary Figure 2

Supplementary Figure 2

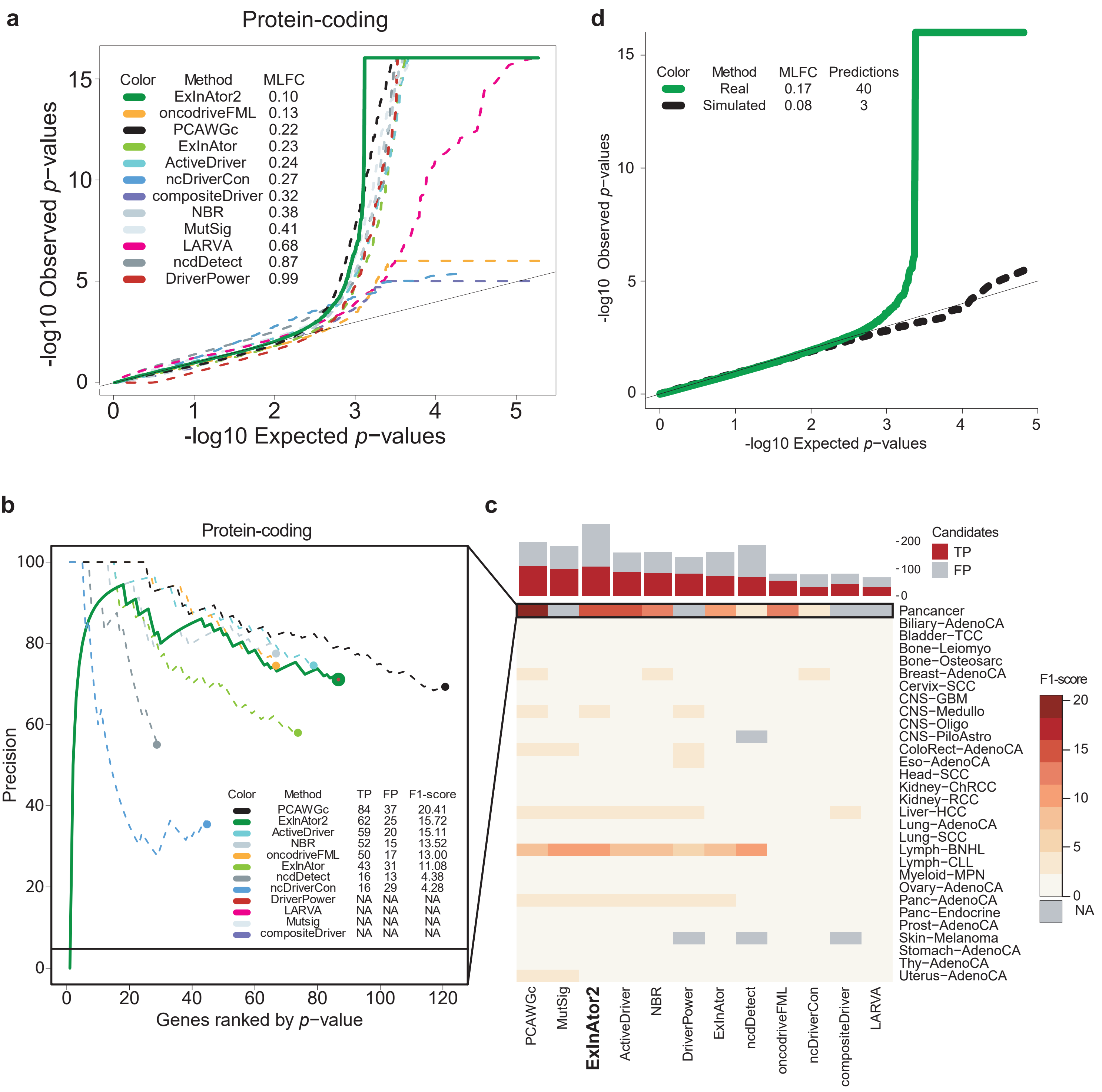

### Supplementary Figure 3

Supplementary Figure 3

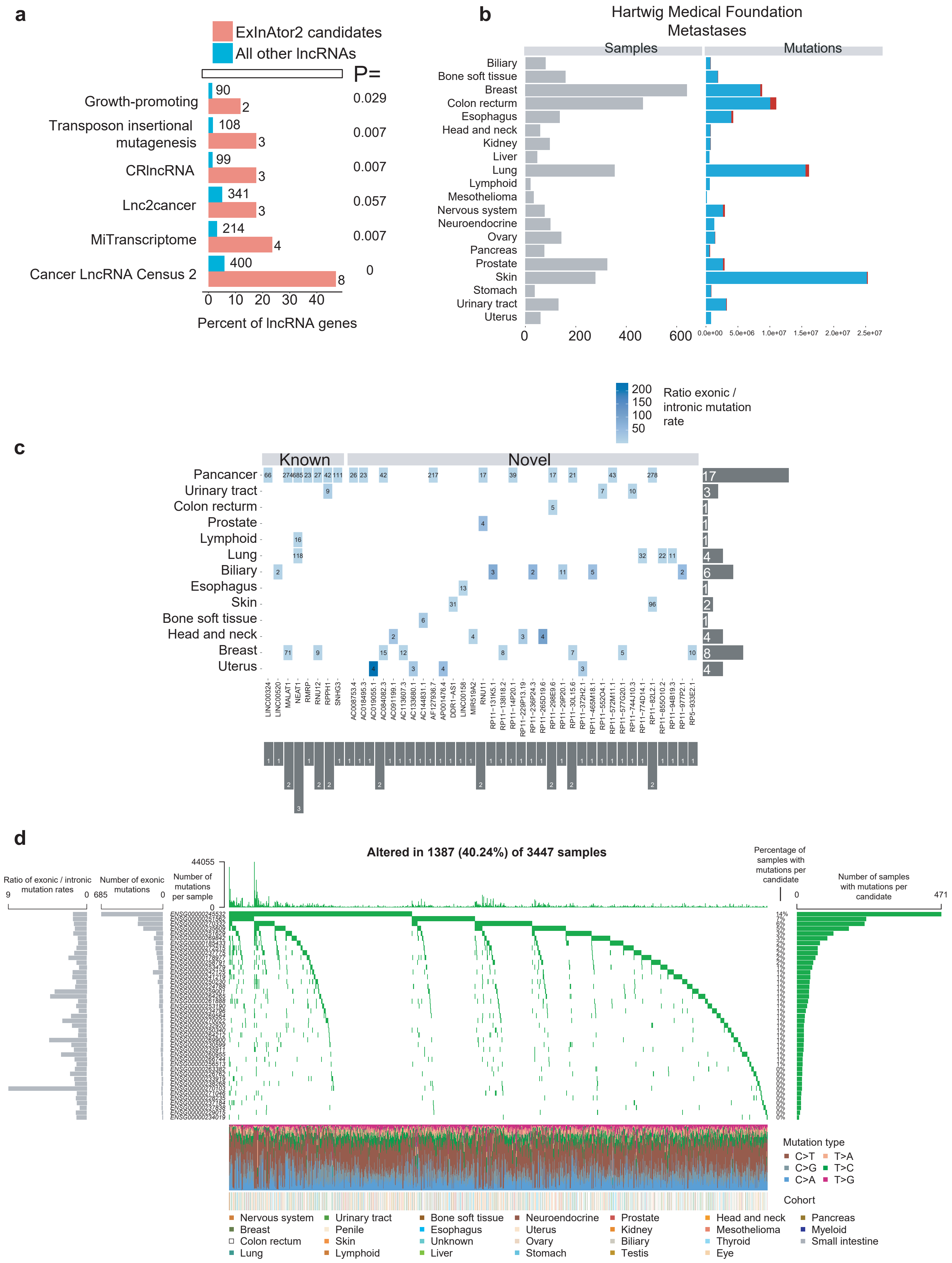

### Supplementary Figure 4

Supplementary Figure 4

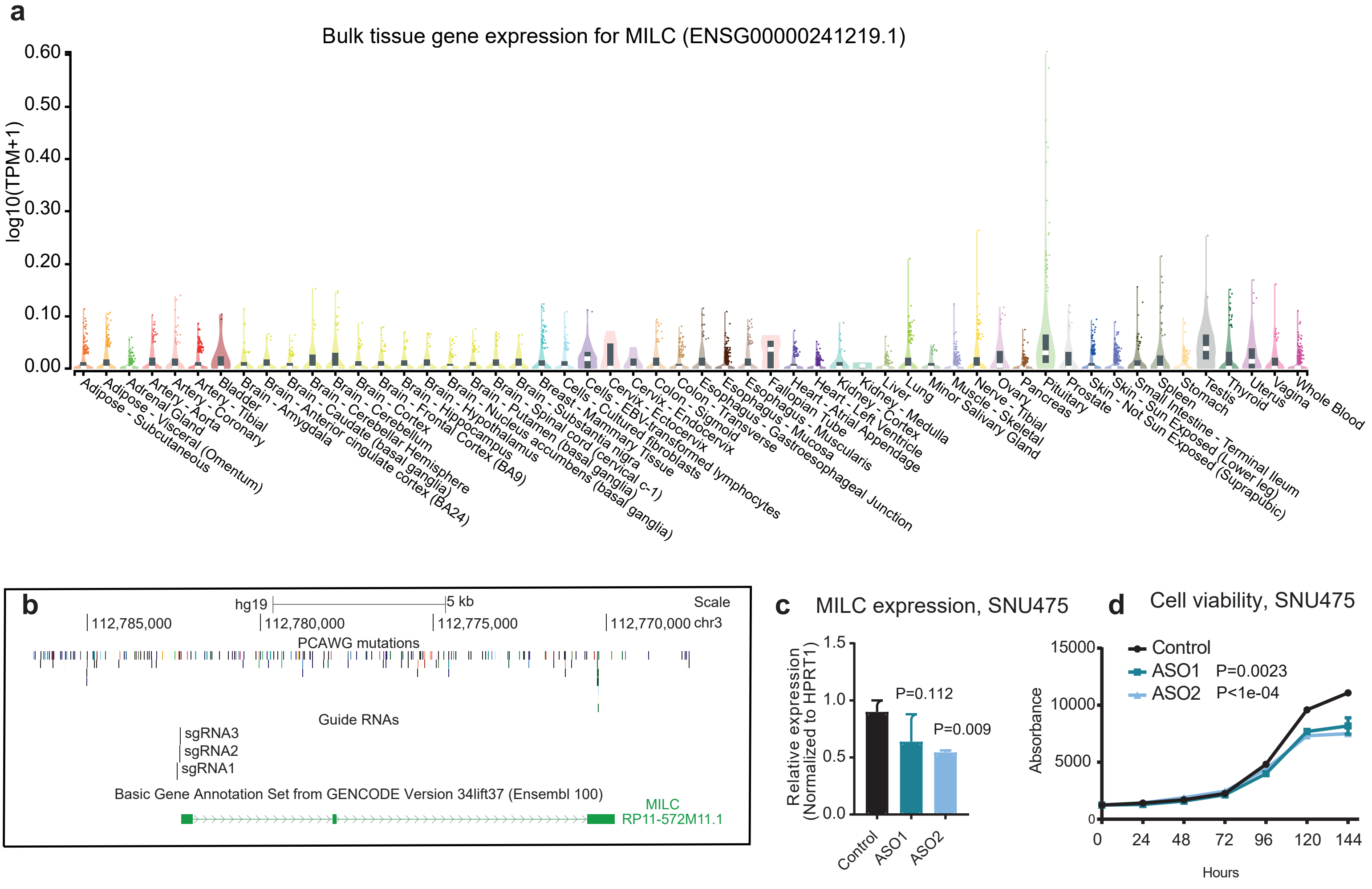

### Supplementary Figure 5

#### Supplementary Figure 5

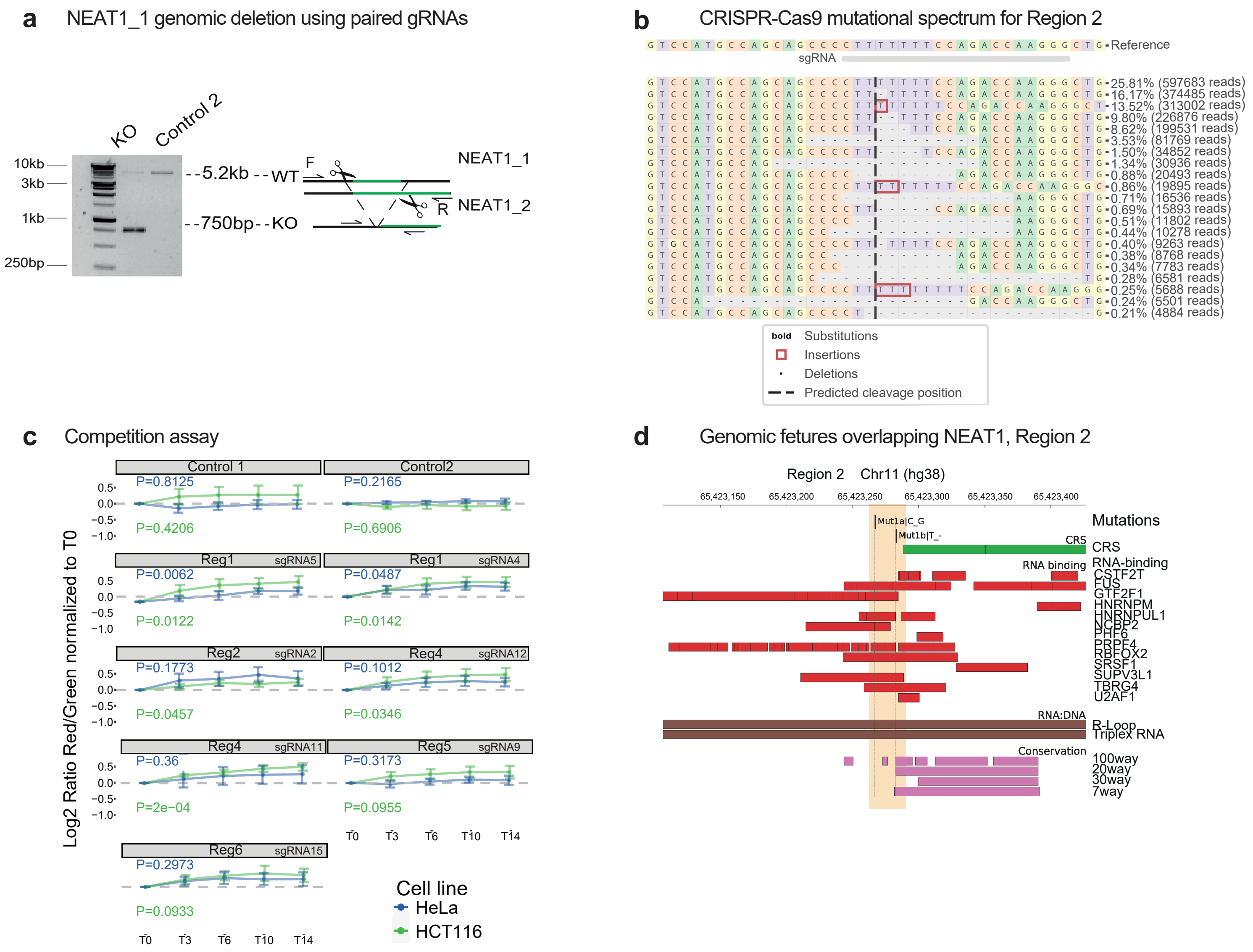
